# DNA double-strand breaks induced by reactive oxygen species promote DNA polymerase IV activity in *Escherichia coli*

**DOI:** 10.1101/533422

**Authors:** Sarah S. Henrikus, Camille Henry, John P. McDonald, Yvonne Hellmich, Steven T. Bruckbauer, Megan E. Cherry, Elizabeth A. Wood, Roger Woodgate, Michael M. Cox, Antoine M. van Oijen, Harshad Ghodke, Andrew Robinson

## Abstract

Under many conditions the killing of bacterial cells by antibiotics is potentiated by damage induced by reactive oxygen species (ROS). In most bacteria, ROS primarily target biomolecules such as proteins and DNA. Damage to DNA, particularly in the form of double-strand breaks (DSBs), is a major contributor to cell death. DNA polymerase IV (pol IV), an error-prone DNA polymerase produced at elevated levels in cells experiencing DNA damage, has been implicated both in ROS-dependent killing and in DSB repair (DSBR). Here, we show using single-molecule fluorescence microscopy that ROS-induced DSBs promote pol IV activity in two ways. First, exposure to the DNA-damaging antibiotics ciprofloxacin and trimethoprim triggers an SOS-mediated increase in intracellular pol IV concentration that is strongly dependent on both ROS and DSBR. Second, in cells that constitutively express pol IV, co-treatment with a ROS mitigator dramatically reduces the number of DSBs as well as pol IV foci formed, indicating a role of pol IV in the repair of ROS-induced DSBs.

**Significance:** Many antibiotics induce an accumulation of reactive oxygen species (ROS) in bacterial cells. ROS-induced damage to DNA, in particular formation of double-strand breaks (DSBs), potentiates killing by several bactericidal antibiotics. Here we used single-molecule fluorescence microscopy to reveal new links between ROS-induced DSBs and the activity of error-prone DNA polymerase IV (pol IV). We found that antibiotic-induced up-regulation of pol IV production requires active formation of DSB intermediates and can be supressed by ROS mitigators. The formation of pol IV foci, which reflect DNA-binding events, also requires DSB repair. Our findings support a major role for pol IV in DSB intermediates and reveal new details of how antibiotic treatment can potentially drive the development of antibiotic resistance in bacteria.

## Main

Many antibiotics induce the accumulation of reactive oxygen species (ROS) within bacterial cells (1–4). These highly reactive molecules cause widespread damage to biomolecules. It is becoming clear that secondary DNA lesions induced by ROS, such as double-strand breaks (DSBs) (5,6) and oxidized nucleotides (7, 8), potentiate killing by bactericidal antibiotics. This phenomenon of secondary lesion formation, which has been described for several antibiotic classes with different primary modes of action, is known as the common killing mechanism (8–15). A well-studied model of the common killing mechanism is the fluoroquinolone antibiotic ciprofloxacin, a DNA gyrase inhibitor, for which killing is strongly potentiated by ROS accumulation (12). A second well-studied model of the common killing mechanism is trimethoprim (13), an antibiotic that inhibits folic acid production and consequently induces thymineless death (TLD). Recent work indicates that TLD involves the accumulation of ROS, which lead to the formation of DSBs (5).

Two mechanisms for ROS-induced DSB formation have been proposed in *E. coli*. The first invokes oxidization of the cellular nucleotide pool, leading to increased incorporation of oxidized nucleotide triphosphates (e.g. 8-oxo-dGTP) into the DNA, for instance, by DNA polymerase IV (7, 16). Subsequent initiation of base-excision repair (BER) creates single-stranded DNA (ssDNA) gaps. In cases where BER is initiated at nearby sites, DSBs may be formed (7,15,16). Evidence for a second mechanism of ROS-dependent DSB formation has emerged from a recent mechanistic study of TLD in *Escherichia coli* (5, 13). The ROS-driven potentiation of killing by both antibiotic treatment and TLD can be abrogated through the addition of ROS mitigators to the culture medium (1,5,12). For example, dimethyl sulfoxide (DMSO) and 2,2’-bipyridine (BiP), both, effectively mitigate the accumulation of antibiotic-induced ROS (5, 17). Using microscopy to quantify ssDNA gaps and DSBs in cells undergoing TLD, Hong and co-workers discovered that thymine starvation initially leads to the accumulation of ssDNA gaps, which are subsequently converted to DSBs in an ROS-dependent process (5). In cells treated with ROS mitigators, gaps were not converted to DSBs and thymine starvation was largely abolished (5). For ciprofloxacin, a DNA gyrase inhibitor, a second, ROS-independent pathway exists in which gyrase-stabilized cleavage complexes dissociate, creating a DSB directly (9–11,18).

Several lines of evidence implicate pol IV in ROS-dependent DSB formation and processing. Pol IV efficiently incorporates 8-oxo-dGTP into DNA *in vitro* (7). Cells over-expressing pol IV exhibit ROS-dependent lethality (7,16,19). Similarly, cells lacking pol IV and pol V are partially protected against killing by ampicillin under conditions where ROS concentrations are increased (7). These observations suggest that pol IV promotes the formation of DBSs due to the BER-mediated removal of closely spaced 8-oxo-dGTPs incorporated by pol IV (16). Other studies indicate that pol IV has a role in the repair of DSBs (20,21,30,31,22–29): First, pol IV physically interacts with the RecA recombinase and RecA nucleoprotein filaments (RecA*); a key player in DSB repair (DSBR) (26, 32). This interaction might facilitate pol IV to function in strand exchange (33). Second, fluorescently labelled pol IV colocalizes with RecA extensively at sites of induced DSBs when expressed from a low-copy plasmid (27). Similarly, in cells treated with ciprofloxacin, pol IV highly colocalizes with RecA* structures (32). Third, genetic studies reveal that the gene encoding pol IV, *dinB*, is required for both induced and spontaneous error-prone DSBR (20–25). Fourth, intermediates of DSBR known as recombination D-loops are efficiently utilized as substrates by pol IV *in vitro* (28, 34).

Interestingly, the mutagenic potential of pol IV is modulated by UmuD and the recombinase RecA (26,29–31). UmuD induces error-free synthesis of pol IV (26), promoting long-lived association of pol IV with the DNA (32). Following UmuD cleavage, pol IV however operates error-prone (26) and pol IV association with DNA is inhibited (32). Furthermore, pol IV operates in an error-prone manner in recombination intermediates *in vitro* (29). Error-prone activity of pol IV in recombination intermediates might be induced due to the interaction of pol IV with RecA (26, 29). Beyond this, RecA promotes DNA synthesis by pol IV in replisomes *in vitro* (30). In the presence of RecA, pol IV can also bypass alkylation lesions more efficiently (31). In addition, RecA nucleoprotein formation on single-stranded DNA is a major trigger for SOS induction and thus increased pol IV expression (35). For some antibiotics, it has however been shown that the SOS response is mostly triggered following DSB processing by RecBCD (36, 37). Notably, upon induction of the SOS response, the cellular concentration of pol IV increases significantly (38, 39). Despite these observations, it remains unclear if pol IV primarily works in recombination intermediates or in the context of replisomes in cells.

Here, we used single-molecule fluorescence microscopy to investigate whether ROS, and ROS-mediated DSBs, influence pol IV expression and association with the nucleoid in cells. We used two antibiotics which alter DNA replication and for which killing is known to involve ROS generation; ciprofloxacin and trimethoprim (5, 12). We further showed that DSB resection is necessary for the formation of pol IV foci, even in cells expressing high concentrations of pol IV (constitutive SOS, *lexA51* mutants, here: *lexA*[Def] mutants), suggesting that pol IV mainly operates on recombination intermediates.

## Results

### ROS potentiate the expression levels and activity of pol IV

We set out to investigate the influence of antibiotic-induced ROS on pol IV activity by monitoring fluorescently-tagged single pol IV molecules in cells. Toward that objective, we first compared pol IV expression levels and its dynamic behavior under normal conditions (no DMSO) and ROS-mitigating conditions (DMSO added) in response to antibiotic treatment. Cells were treated with i) ciprofloxacin alone, ii) ciprofloxacin and DMSO in combination, iii) trimethoprim alone or iv) trimethoprim and DMSO in combination (Fig. 1, *SI Appendix*, Fig. 1*A*).

**Figure 1.**
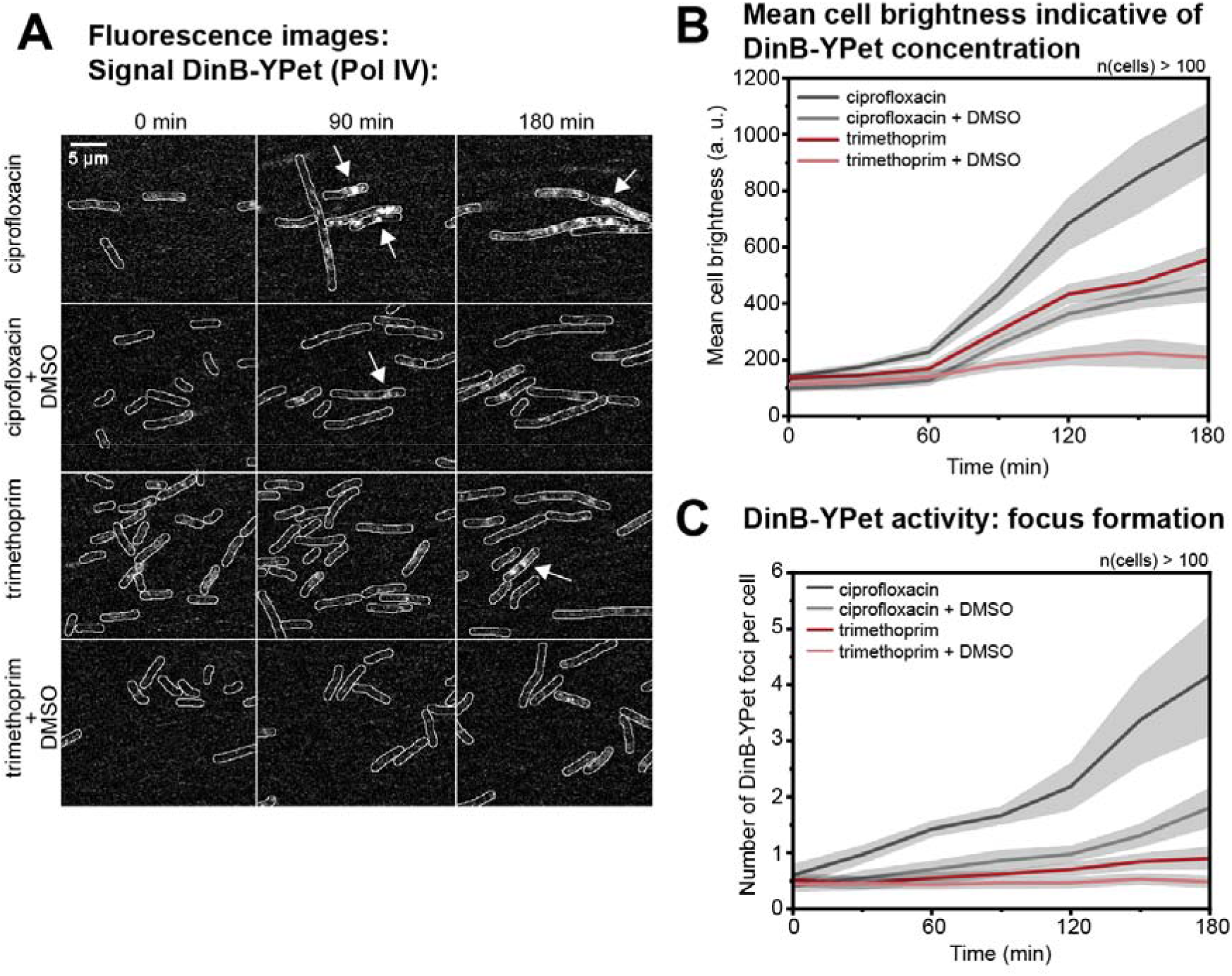
Pol IV concentration and activity following ciprofloxacin or trimethoprim treatment under normal conditions or ROS-mitigating conditions. (*A*) Fluorescence images showing cells expressing DinB-YPet (Pol IV) at 0, 90 and 180 min (left to right) after ciprofloxacin-alone, ciprofloxacin-DMSO, trimethoprim-alone or trimethoprim-DMSO treatment (top to bottom). Scale bar represents 5 µm. (*B*) Concentration of DinB-YPet during stress. Mean cell brightness is plotted against time (ciprofloxacin-alone: dark grey line, ciprofloxacin-DMSO: light grey line, trimethoprim-alone: magenta line, trimethoprim-DMSO: light magenta line). At each time-point, data are derived from >100 cells. Grey shaded error bands represent standard error of the mean. (*C*) Number of DinB-YPet foci per cell are plotted against time (ciprofloxacin-alone: dark grey line, ciprofloxacin-DMSO: light grey line, trimethoprim-alone: red line, trimethoprim-DMSO: light red line). At each time-point, data are derived from >100 cells. Grey shaded error bands represent standard error of the mean.

Prior to live-cell imaging, we first established that cells expressing fluorescent protein fusions of DinB, τ (replisome marker) and UmuC (component of DNA polymerase V, pol V) exhibited wild-type oxidative stress responses upon antibiotic treatment (ciprofloxacin or trimethoprim) administered either alone or along with the ROS mitigator (DMSO) (*SI Appendix*, Fig. 2). In the presence of ROS, *E. coli* cells induce the peroxide and/or superoxide stress responses in which expression of superoxide dismutase, alkyl hydroperoxidase and Fe^3+^ enterobactin transporter genes are upregulated (reviewed in (3,4,40–44)). Therefore, we developed an assay to monitor expression of *gfp* from ROS-regulated and iron-responsive promoters in cells treated with ciprofloxacin, trimethoprim or hydrogen peroxide (as a control). We further tested if the addition of DMSO suppressed the accumulation of ROS (*SI Appendix*, Fig. 3, 4, 5). For this purpose, we constructed three plasmids that express GFP (fast-folding GFP, *sf-gfp* (45)) from the ROS-regulated promoters of *sodA* (notably regulated by superoxides/redox active compound *via* SoxRS and by the iron (Fe^2+^) concentration via Fur (4, 41), *SI Appendix*, Fig. 3*A*), *ahpC* (regulated by OxyR (4,42,43), *SI Appendix*, Fig. 4*A*) or *fepD* (regulated by Fur pathway; iron homeostasis (44), *SI Appendix*, Fig. 5*A*). Following hydrogen peroxide treatment, the addition of DMSO reduced the expression of the GFP reporter from the plasmid-based *sodA* and *fepD* promoters by 30% (30 mM hydrogen peroxide at t = 8 h, *SI Appendix*, Fig. 3*C*, 4*D*). This reduction in GFP signal is not due to DMSO quenching fluorescence (*SI Appendix*, Fig. 6). Increased expression from the *ahpC* promotor was delayed by ∼3 h (30 and 100 mM hydrogen peroxide, *SI Appendix*, Fig. 4*D*). For ciprofloxacin-treated cells, the addition of DMSO reduced expression from the *fepD* promotor by 50% (5, 10, 20 and 40 ng/mL at t = 8 h, *SI Appendix*, Fig. 5*B*). For trimethoprim-treated cells, the addition of DMSO reduced the expression from the *ahpC* and *fepD* promotors by 50% (0.1 and 0.3 μg/mL at t = 8 h, *SI Appendix*, Fig. 4*B*, 5*B*). Together, these results indicate that (i) ciprofloxacin and trimethoprim generate ROS in cells (consistent with previous work (5, 46)) and (ii) DMSO reduced the expression from ROS-sensitive promoters, following hydrogen peroxide, ciprofloxacin and trimethoprim treatment, implying that ROS levels were effectively reduced by the addition of DMSO.

Following antibiotic addition, we recorded time-lapse movies capturing fluorescence from *Escherichia coli* cells expressing a functional, YPet fusion of the DinB gene from its native promoter (*SI Appendix*, Fig. 1*B, C*, Materials and Methods) (39, 47). We then monitored pol IV concentrations by measuring the fluorescence intensity of DinB-YPet within cells in the presence or absence of DMSO (2% v/v) and monitored DNA binding activities by counting the number of pol IV foci per cell. Treatment with ciprofloxacin resulted in cell filamentation accompanied by a clear increase in DinB-YPet intensity, indicating an increase in the intracellular DinB-YPet concentration (seven-fold increase from 140 to 990 DinB-YPet fluorescence, Fig. 1*A*, *B*; *SI Appendix*, Fig. 7). In a previous study (39), following ciprofloxacin treatment, cells exhibited a similar increase in DinB-YPet concentration; an increase in intracellular DinB-YPet (pol IV) concentrations was measured from 6 ± 1 nM prior to treatment (standard error of the mean, SE) to 34 ± 3 nM (SE) 180 min after ciprofloxacin addition. Interestingly, in this present study, we showed that inclusion of DMSO led to a significant reduction in the expression level of DinB-YPet in ciprofloxacin-treated cells. DMSO was added at the concentration previously tested (*SI Appendix*, Fig. 3, 4, 5). 180 min after ciprofloxacin addition, cellular DinB-YPet intensities were only four-fold higher than basal levels (intensity increase from 100 to 454, Fig. 1*A*, *B*). This final intensity corresponds to a concentration of DinB-YPet equalling 19 ± 2 nM (SE, see *Materials and Methods*), corresponding to a reduction of about 15 nM of ciprofloxacin-induced pol IV. Treatment with trimethoprim alone led to a significant increase in DinB-YPet fluorescence; 180 min after trimethoprim addition, the mean fluorescence intensity increased by more than four-fold (fluorescence intensity increase from 135 to 557, Fig. 1*A*, *B*), corresponding to a final intracellular pol IV concentration of 23 ± 2 nM. Inclusion of DMSO led to a significant reduction in trimethoprim-induced pol IV up-regulation; cellular DinB-YPet fluorescence intensities increased only slightly from 113 to 209, corresponding to a final pol IV concentration of 9 ± 2 nM. Thus, for both antibiotics, addition of DMSO resulted in a significant reduction in the steady state levels of pol IV in response to treatment.

Cells exhibit distinct pol IV foci when individual DinB-YPet molecules bind to DNA and thus experience decreased diffusional mobility (48). Since cells expressing fluorescently tagged catalytically dead pol IV molecules do not exhibit foci (39), the foci observed in response to antibiotic treatment represent pol IV molecules engaged in catalytic functions. Prior to the addition of ciprofloxacin, cells contained on average 0.6 ± 0.2 foci per cell (SE) in the absence of DMSO, and 0.4 ± 0.1 foci per cell in the presence of DMSO (Fig. 1*C*). Following treatment with ciprofloxacin alone, the number of foci steadily increased. By 180 min, cells had 4.2 ± 1.1 foci per cell. Upon ciprofloxacin-DMSO treatment, cells contained 1.8 ± 0.4 foci per cell; a > 50% reduction compared to ciprofloxacin-alone measurements. Prior to the addition of trimethoprim, cells contained on average 0.5 ± 0.1 foci (SE) in the absence of DMSO and 0.4 ± 0.1 foci in the presence of DMSO. Trimethoprim-alone treatment induced a slight increase in the number of DinB-YPet foci with 0.9 ± 0.2 per cell (SE) at 180 min. This is lower than the number of foci observed for ciprofloxacin-DMSO treatment (1.8 ± 0.4 per cell), despite the measured pol IV concentration being marginally higher after trimethoprim-alone treatment (Fig. 1*C*). Strikingly, cells treated with both trimethoprim and DMSO did not show any increase in DinB-YPet foci after trimethoprim addition (0.5 ± 0.1 foci per cell at 180 min; Fig. 1*C*). Together, these results demonstrate that for cells treated with ciprofloxacin or trimethoprim, addition of DMSO supresses the drug-induced increases in DinB-YPet concentration, as well as the binding of pol IV to DNA, as evidenced by a reduction in the number of DinB-YPet foci. Importantly, the concentration of pol IV and its extent of DNA-binding are not directly correlated as the trimethoprim-alone and ciprofloxacin-DMSO treatments induced similar DinB-YPet concentrations, but different numbers of DinB-YPet foci.

### ROS-induced double-strand breaks trigger the SOS response

Reasoning that the decreased induction of *dinB-YPet* expression in cells co-treated with DMSO likely resulted from attenuation of the SOS response, we repeated the time-lapse experiments (*SI Appendix*, Fig. 1*B, C*) on cells that carried an SOS-reporter plasmid, in which GFP is expressed from the SOS-inducible *sulA* promoter (pUA66 P*_sulA_*-*gfp*; fast-folding GFP, *gfpmut2* (49)). In the absence of any antibiotic treatment, cells exhibit very low fluorescence intensity, consistent with the repression of the *sulA* promoter in the absence of exogenously applied DNA damage (Fig. 2*A*, ‘0 min’). SOS levels were similarly low for cells grown in the presence of DMSO. Cells exhibited robust SOS induction upon treatment with ciprofloxacin as evidenced by the increase in GFP fluorescence in the 180 min time window after addition of ciprofloxacin (170 fold induction, Fig. 2*B*). Consistent with our hypothesis, SOS induction was strongly inhibited upon inclusion of DMSO during ciprofloxacin treatment (13 fold induction at 180 min; Fig. 2*B*). A similar reduction in ROS and SOS levels has been observed in cells following co-administration of ciprofloxacin with another ROS mitigator, *N*-acetylcysteine (50). Cells exposed to trimethoprim exhibited a delay in SOS induction, however, even in this case, high levels of SOS induction (100 fold induction) were supressed by the addition of DMSO (2 fold induction for combined treatment with trimethoprim and DMSO; Fig. 2*B*). Notably, the addition of a different ROS mitigator, 2,2’-bipyridine (BiP, 0.35 mM, 0.5 x MIC (5)), similarly supressed the induction of the SOS response (Fig. 2). These results were also confirmed using plate-reader assays (*SI Appendix*, Fig. 8).

**Figure 2.**
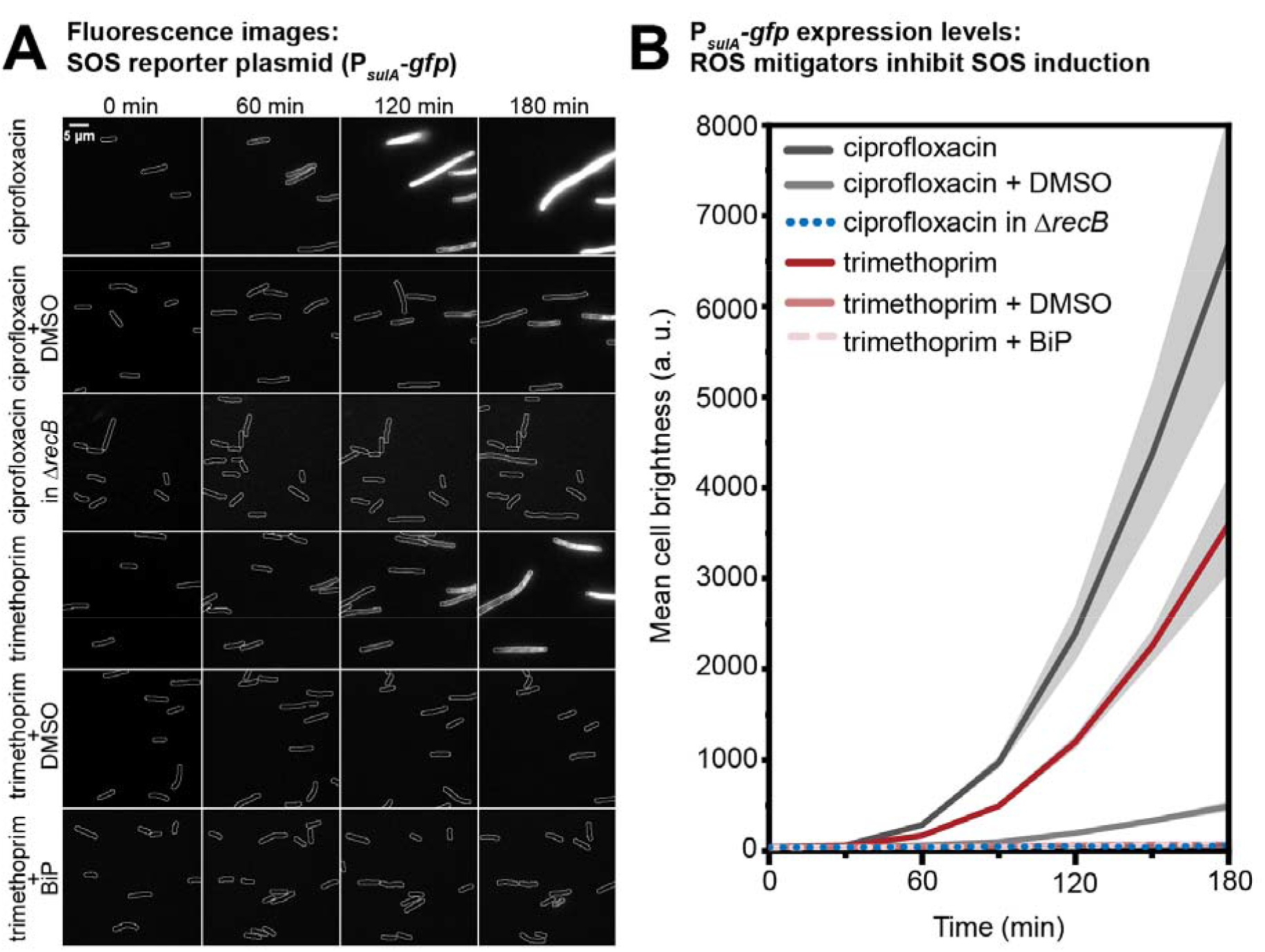
P*_sulA_-gfp* expression levels (SOS response levels) following ciprofloxacin or trimethoprim treatment under normal or ROS-mitigating conditions in different genetic backgrounds. (*A*) Fluorescence images showing the expression of GFP from a SOS reporter plasmid (P*_sulA_*-*gfp*) at 0, 60, 120 and 180 min (left to right) after ciprofloxacin-alone, ciprofloxacin-DMSO, ciprofloxacin-alone in Δ*recB*, trimethoprim-alone, trimethoprim-DMSO or trimethoprim-BiP treatment (top to bottom). Scale bar represents 5 µm. (*B*) GFP expression levels from the *sulA* promotor during stress. Mean cell intensity is plotted against time (ciprofloxacin-alone: dark grey line, ciprofloxacin-DMSO: light grey line, ciprofloxacin in Δ*recB*: purple, dotted line, trimethoprim-alone: red line, trimethoprim-DMSO: light red line, trimethoprim-BiP: rose-colored, dashed line). At each time-point, data are derived from >100 cells. Grey shaded error bands represent standard error of the mean.

We reasoned that the suppression of SOS by ROS mitigators might reflect a reduction in the formation and processing of DSBs. Cells lacking *recB* fail to induce SOS upon treatment with nalidixic acid, suggesting that end-resection products formed by RecBCD might be sites of SOS induction (36, 37). Since ciprofloxacin and nalidixic acid both target DNA gyrase (9,11,51), we repeated the GFP reporter measurements in cells lacking *recB* (SSH111, Δ*recB* P*_sulA_*-*gfp*) to determine if SOS induction by ciprofloxacin is also dependent on DSB processing. The deletion of *recB* strongly inhibited the SOS response following ciprofloxacin treatment (0.4 fold induction at 180 min in comparison to *recB^+^*, Fig. 2, *SI Appendix,* Fig. 9). While *recB* deletions are known to reduce survival in cells treated with ciprofloxacin (52), we observed that most cells lacking *recB* continued to grow and divide during the 180 min time-lapse measurement (*SI Appendix*, Fig. 7, 9), indicating that the lack of SOS induction observed for ciprofloxacin-treated *recB*-deficient cells did not stem from gross inhibition of all cellular functions. Plate reader assays did not reveal a sustained increase in cell mass for *recB* deletion cells following ciprofloxacin treatment (*SI Appendix*, Fig. 8*A*, last column), suggesting that the initial growth observed by microscopy stagnates soon after the 180 min observation window.

To more directly investigate if ROS create DSBs following ciprofloxacin and trimethoprim treatment, we imaged cells expressing a fluorescent fusion of the DSB reporter MuGam (53) to the photoactivatable mCherry protein (PAmCherry1 (54), *SI Appendix*, Fig. 1A, *C*). MuGam-PAmCherry was expressed from a plasmid (Fig. 3*A*). For these single-molecule microscopy experiments, expression of MuGam was induced using 0.003% L-arabinose at MuGam expression levels that had minimal effects on survival upon drug treatment (*SI Appendix*, Fig. 10). In the absence of antibiotic, cells exhibited 0.3 ± 0.1 MuGam foci per cell with 74% of cells containing no foci (Fig. 3b, c). Two hours after ciprofloxacin treatment, cells contained increased number of MuGam foci per cell (4.9 ± 0.3 foci with 1.6% of cells containing no foci, Fig. 3*C*). Consistent with DMSO mitigating ROS, DMSO addition reduced the number of MuGam foci per cell (2.2 ± 0.2 foci with 21% of cells containing no foci, Fig. 3*C*), indicating a significant contribution of ROS to the formation of DSBs during ciprofloxacin treatment. In agreement with a previous study (5), we observed that trimethoprim treatment generates DSBs (1.9 ± 0.1 MuGam foci with 22% of cells containing no foci, Fig. 3*C*). These DSBs are ROS-induced as the addition of DMSO prevents the formation of these DSBs (0.5 ± 0.1 foci with 59% of cells containing no foci, Fig. 3*C*). In contrast, in a recent study using sub-inhibitory concentrations of ciprofloxacin, reactive oxygen species do not induce additional DSBs (55).

**Figure 3.**
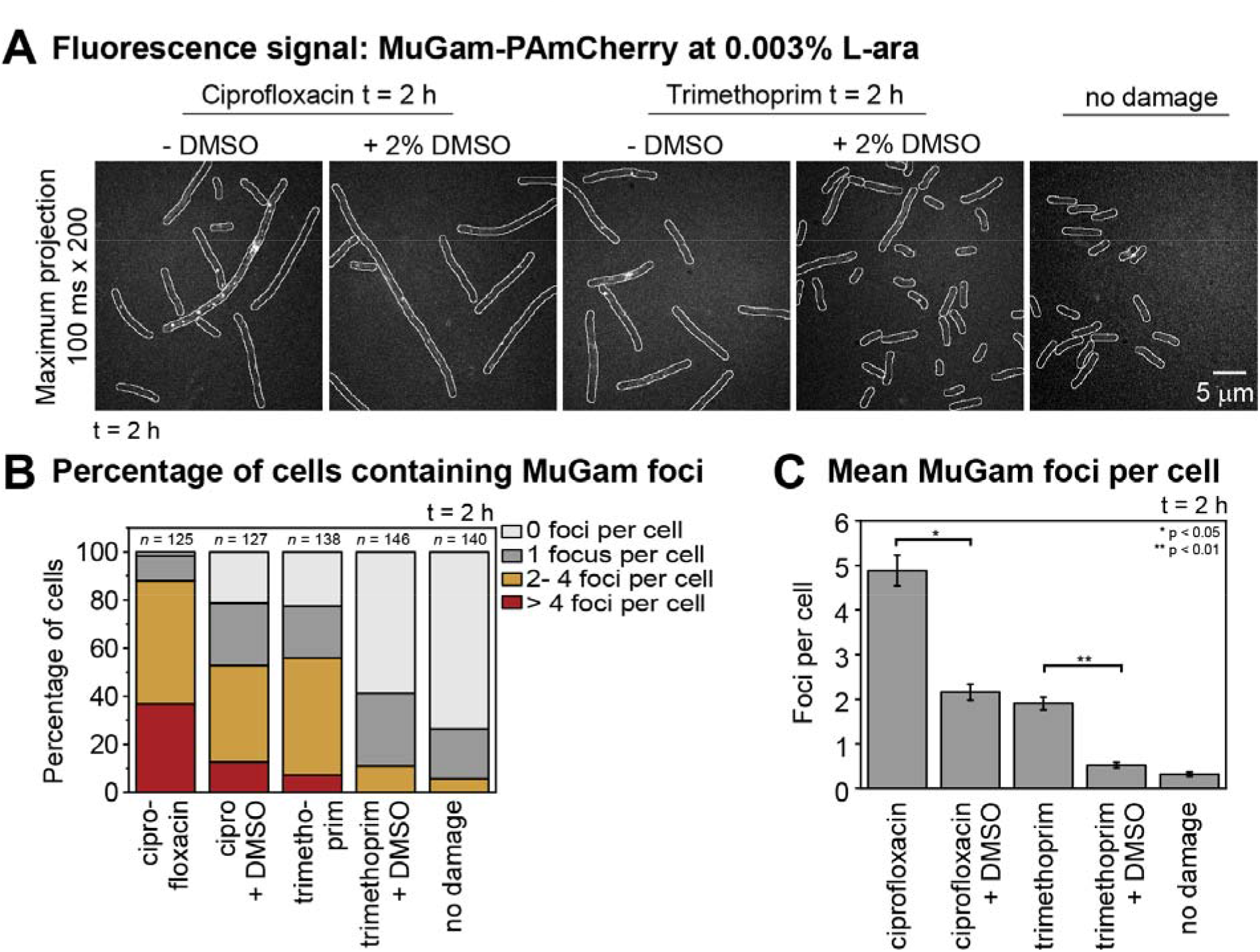
Number of MuGam-PAmCherry foci per cell following ciprofloxacin or trimethoprim treatment under normal conditions or ROS-mitigating conditions in different genetic backgrounds. (*A*) Fluorescence signal from MuGam-PAmCherry at 0.003% L-arabinose: Maximum projections over 100 ms x 200 frames showing MuGam-PAmCherry foci. From left to right: MuGam signal after 2 h treatment with ciprofloxacin, ciprofloxacin + DMSO, trimethoprim, trimethoprim + DMSO, no damage. (*B*) Percentage of cells containing MuGam foci: 0 foci (light grey), 1 focus (grey), 2-4 foci (amber) and > 4 foci (red). Cells were treated with ciprofloxacin (*n* = 125), ciprofloxacin + DMSO (*n* = 127), trimethoprim (*n* = 138), trimethoprim + DMSO (*n* = 146), or experienced no damage (*n* = 140). (*C*) Mean number of MuGam foci per cell. Cells were treated with ciprofloxacin (*n* = 125), ciprofloxacin + DMSO (*n* = 127), trimethoprim (*n* = 138), trimethoprim + DMSO (*n* = 146), or experienced no damage (*n* = 140). The error bars represent standard error of the mean over the number of cells. * for p < 0.05; ** for p < 0.01.

Taken together our measurements indicate that antibiotic-induced ROS generate DSBs and potentiate the SOS response. Furthermore, SOS induction levels are dependent on *recB* DSB processing in cells treated with ciprofloxacin. Together the results are consistent with a model in which the SOS response is triggered or potentiated in antibiotic-treated cells via ROS-induced DSBs, leading to increased levels of pol IV in cells.

### Double-strand break resection creates substrates for pol IV

Having established conditions under which ROS create a majority of DSBs in cells as well as binding sites for pol IV upon antibiotic treatment, we next set out to characterize pol IV behavior during DSBR in response to antibiotic treatment. To that end, we tested if pol IV primarily forms foci following DSB resection, suggestive of pol IV having a major role in DSBR. Therefore, we examined the extent of DinB-YPet focus formation in ciprofloxacin and trimethoprim treated cells, comparing backgrounds that permitted (*recB*^+^) or prevented (Δ*recB*) DSB processing. Additionally, we monitored the formation of DinB-YPet foci while using DMSO to modulate the number of antibiotic-induced DSBs (Fig. 3). To separate effects on focus formation from effects on DinB-YPet expression, these measurements were carried out in a *lexA*(Def) background (56) (*dinB-YPet dnaX-mKate2 lexA*[Def]). These cells constitutively express DinB-YPet at levels consistent with SOS induced levels, even in the absence of DNA damage (39). To capture DinB-YPet binding events on the time-scale of seconds, we recorded burst acquisitions of the DinB-YPet signal (300 x 50 ms exposures taken every 100 ms, *SI Appendix*, Fig. 1*A, D*).

Consistent with the results from our previous study (39), close to zero DinB-YPet foci were observed in *lexA*(Def) cells in the absence of antibiotic (0.08 ± 0.05 foci per cell, Fig. 4). In contrast, *lexA*(Def) cells treated with ciprofloxacin for 60 min exhibited clear foci (1.83 ± 0.15 foci per cell, Fig. 4*B*, *C*). Co-treatment with ciprofloxacin and DMSO yielded fewer foci (1.02 ± 0.13 foci per cell, Fig. 4 *B*, *C*). The deletion of *recB* resulted in a striking loss of DinB-YPet foci (0.23 ± 0.05 foci per cell, Fig. 4 *B*, *C*). *lexA*(Def) cells treated with trimethoprim for 60 min contained multiple DinB-YPet foci (2.6 ± 0.18 foci per cell), whereas cells treated with both trimethoprim and DMSO contained few foci (0.19 ± 0.06). Trimethoprim-treated Δ*recB* cells also contained very few foci (0.14 ± 0.05). Similar effects were observed in *lexA*^+^ cells, although reductions in focus formation were conflated with reductions in DinB-YPet expression levels (Fig. 1*C*). Taken together these results demonstrate that pol IV is normally active at ROS-induced, RecBCD-processed DSBs in cells treated with ciprofloxacin or trimethoprim. Consistent with this, we have demonstrated that pol IV co-localizes with RecA* features in cells treated with ciprofloxacin (32).

**Figure 4.**
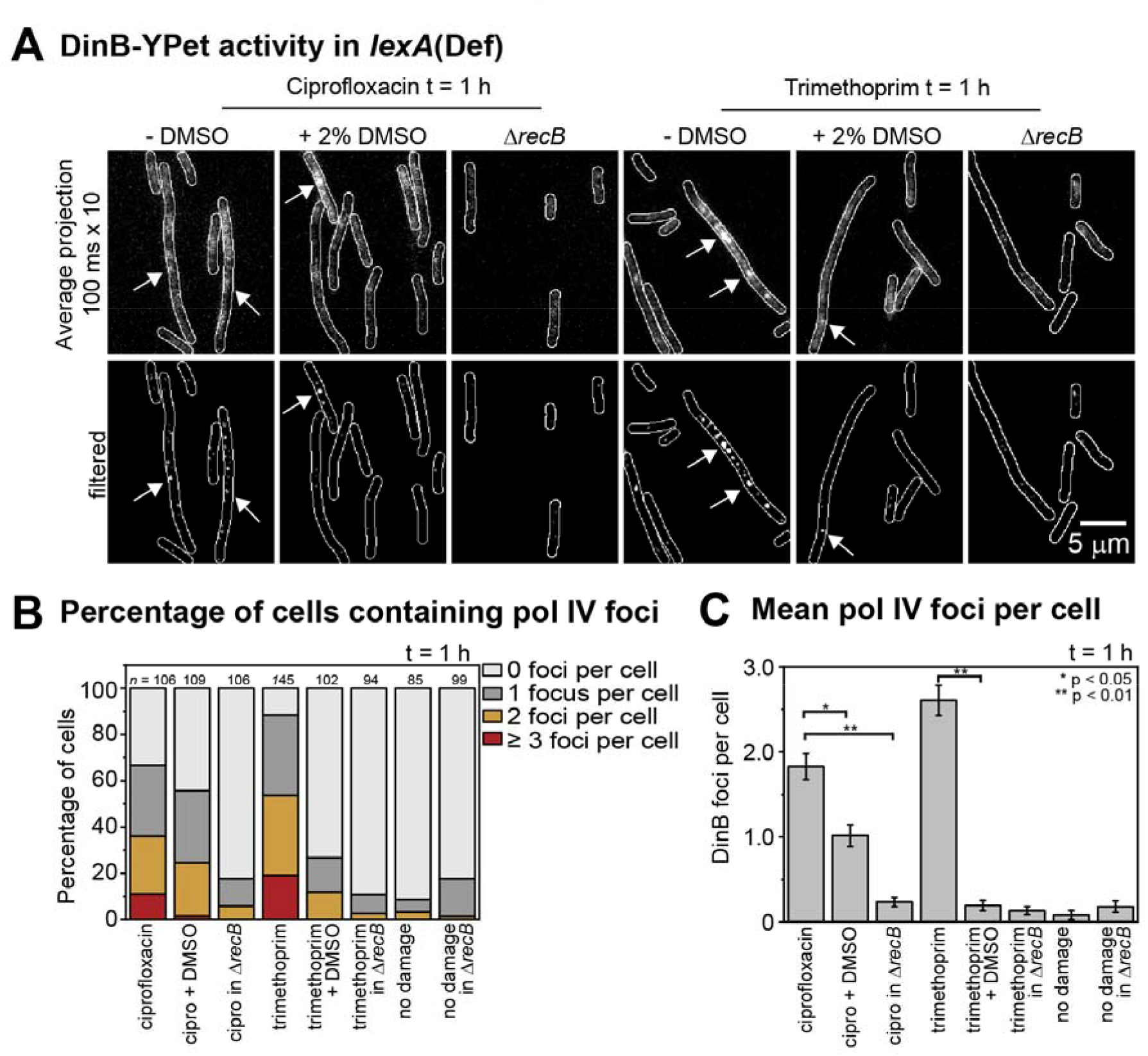
Number of pol IV foci per cell in *lexA*(Def) cells following ciprofloxacin or trimethoprim treatment under normal conditions or ROS-mitigating conditions. (*A*) Upper row: Average projection in time (100 ms x 10 frames) showing DinB-YPet (pol IV) foci. Bottom row: Discoidal filtered projections. Cells were treated for 60 min prior to imaging. (*B*) Percentage of cells containing pol IV foci: 0 foci (light grey), 1 focus (grey), 2 foci (amber) and ≥ 3 foci (red). Cells were treated with ciprofloxacin (*n* = 106), ciprofloxacin + DMSO (*n* = 109), ciprofloxacin in Δ*recB* (*n* = 106), trimethoprim (*n* = 145), trimethoprim + DMSO (*n* = 102), trimethoprim in Δ*recB* (*n* = 94) experienced no damage for wild-type (*n* = 85) and Δ*recB* (*n* = 99). (*C*) Number of DinB-YPet foci per cell. Error bars represent standard error of the mean. Number of cells included in analysis: *n*(ciprofloxacin) = 106, *n*(ciprofloxacin-DMSO) = 109, *n*(ciprofloxacin in Δ*recB*) = 106, *n*(trimethoprim) = 145, *n*(trimethoprim-DMSO) = 102, *n*(trimethoprim in Δ*recB*) = 94, *n*(untreated *recB*^+^) = 85, *n*(untreated Δ*recB*) = 99. * for p < 0.05; ** for p < 0.01.

In a previous study (39), we showed that pol IV primarily forms foci away from replisomes, indicating that pol IV has a minor role in facilitating replication restart of stalled replisomes. To investigate if these non-replisomal pol IV foci are ROS-induced, we next determined the percentage of DinB-YPet foci that form in the vicinity of replisomes (fluorescent protein fusion of the pol III _τ_-subunit, _τ_-mKate2). For each experiment, when recording the DinB-YPet signal in *recB*^+^ cells, we also recorded the position of _τ_-mKate2 as in the previous study (39). Ciprofloxacin treatment, which rapidly halts DNA synthesis (57, 58), causes 10% of pol IV foci to bind near replisomes (39). Here we observed that the inclusion of DMSO dramatically increased the relative colocalization of DinB-YPet with replisomes in both *lexA*^+^ and *lexA*(Def) cells treated with ciprofloxacin (*SI Appendix*, Fig. 11, 12). For long-lived pol IV foci (detectable within a 10 s average projection image, Fig. 5*B*, right panel) in the *lexA*(Def) background, 80% of foci colocalized with replisomes under ciprofloxacin-DMSO conditions (Fig. 5*B*). This is consistent with the addition of DMSO having removed the vast majority of non-replisomal substrates for pol IV-dependent DNA synthesis. This observation appears to be consistent with a recent proposal that ROS-mitigation reduces rates of pol IV-dependent mutagenesis (55). For *lexA*(Def) cells treated with trimethoprim, addition of DMSO abolished long-lived pol IV foci entirely (Fig. 5*A*, *C*).

**Figure 5.**
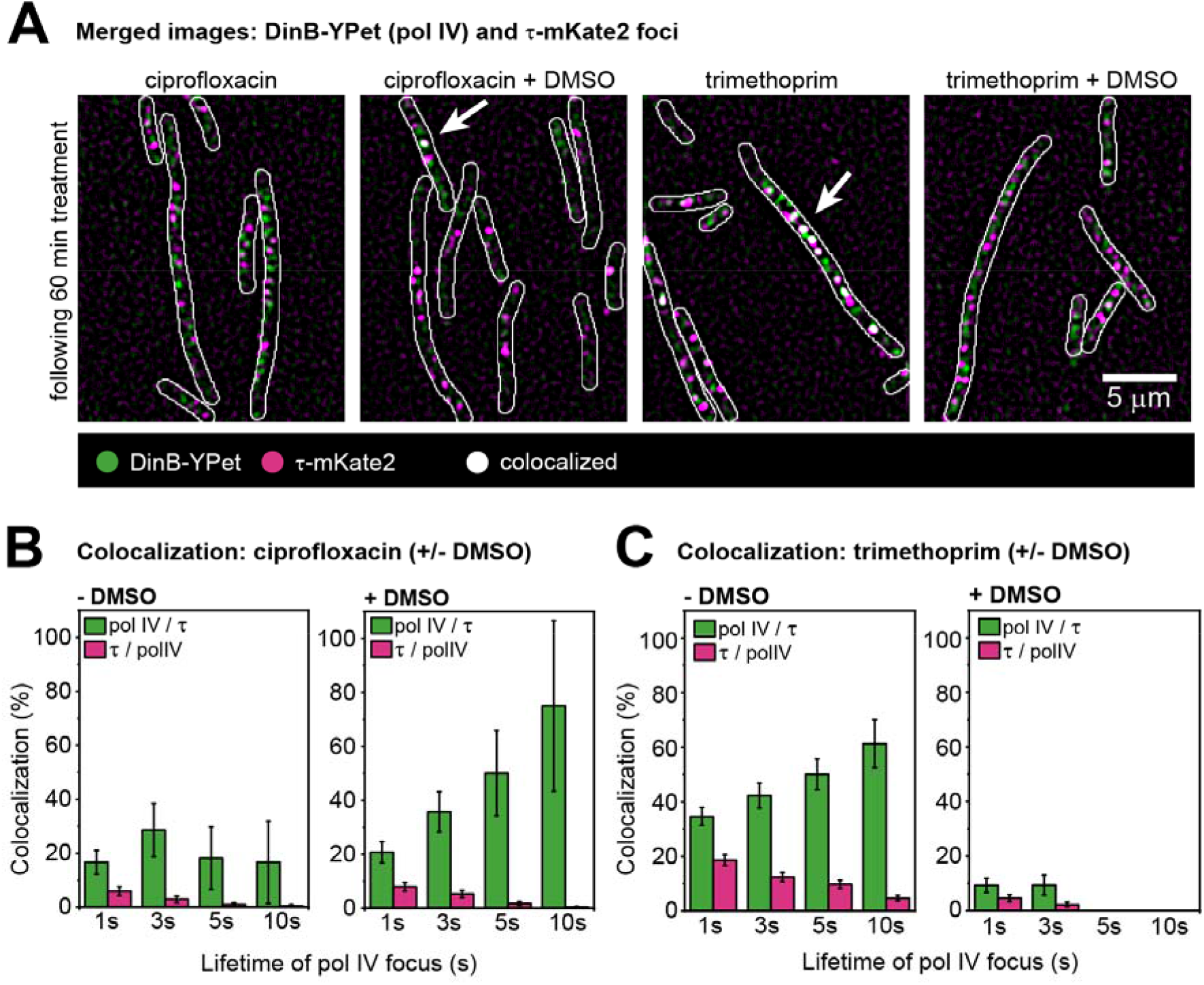
Measuring the colocalization of pol IV and replisomes following ciprofloxacin or trimethoprim treatment ± DMSO in *lexA*(Def) cells. (*A*) DinB-YPet activity at replisomes in *lexA*(Def) cells. Cells were treated for 60 min prior to imaging. Merged images showing DinB-YPet foci in green and τ-mKate2 foci in magenta following ciprofloxacin-alone, ciprofloxacin-DMSO, trimethoprim-alone and trimethoprim-DMSO treatment (from left to right). White arrow points at colocalization event (white focus). Scale bar represents 5 µm. (*B*) Colocalization percentages of pol IV foci that bind at replisomes (green bars) and colocalization percentages of replisomes that contain a pol IV focus (magenta bars) for cells treated with ciprofloxacin-alone (left) or ciprofloxacin-DMSO (right). Colocalization was measured with sets of pol IV foci that last 1, 3, 5 and 10 s. Error bars represent the standard error of the mean. (*C*) Colocalization percentages of pol IV foci that bind at replisomes (green bars) and colocalization percentages of replisomes that contain a pol IV focus (magenta bars) for cells treated with trimethoprim-alone (left) or trimehtoprim-DMSO (right). Colocalization was measured with sets of pol IV foci that last 1, 3, 5 and 10 s. Error bars represent the standard error of the mean.

### ROS do not promote pol V activity

Finally, we explored if ROS-induced DSBs promote a change in the binding activity of the other major error-prone polymerase pol V (UmuDD_2_C) in real time (59). Since pol V is also a member of the SOS regulon (35), we use a *lexA*(Def) background (RW1286, *umuC-mKate2 dnaX-YPet lexA*[Def]) to separate effects on focus formation from effects on UmuC-mKate2 expression.

UmuC foci might form at two stages during the activation of pol V Mut at RecA* filaments and when active pol V Mut complexes synthesize DNA. As before, *lexA*(Def) cells were treated for 60 min with ciprofloxacin-alone, ciprofloxacin-DMSO, trimethoprim-alone or trimethoprim-DMSO (*SI Appendix*, Fig. 1*A*). Burst acquisitions of the UmuC-mKate2 signal were recorded (*SI Appendix*, Fig. 1*D,* 300 x 50 ms exposures taken every 100 ms).

Few UmuC-mKate2 foci were observed in the absence of antibiotic in *lexA*(Def) cells (about 0.32 ± 0.08 foci per cell, *SI Appendix*, Fig. 12). In *lexA*(Def) cells treated with ciprofloxacin or trimethoprim for 60 min, foci were clearly visible (ciprofloxacin: 1.24 ± 0.16 foci per cell; trimethoprim 1.39 ± 0.21 foci per cell). In both cases, co-treatment with DMSO had little effect on the number of UmuC-mKate2 foci (ciprofloxacin-DMSO: 0.99 ± 0.12 foci per cell; trimethoprim-DMSO 1.26 ± 0.16 foci per cell) or on the overall levels of UmuC-mKate2 fluorescence in the cells. Thus in contrast to the effects observed for pol IV, the addition of DMSO had little effect on the formation of UmuC foci. Interestingly, in *lexA*^+^ cells, which express SOS normally, trimethoprim treatment (with or without DMSO) did not lead to the formation of pol V (*SI Appendix*, Fig. 13*A*, *C*). Consistent with this, cleavage of UmuD to UmuD′ was far less efficient in trimethoprim-treated cells than in ciprofloxacin-treated cells (compare *SI Appendix*, Fig. 14*B*, *D*). This suggests that RecA* structures that induce SOS (i.e. increase in the expression levels of SulA and pol IV) may be different from those that mediate the formation of pol V through UmuD cleavage. This result is discussed further below and warrants further investigation.

## Discussion

### ROS-mediated DSBs induce high intracellular concentrations of pol IV

We observed that ROS mitigators reduced levels of SOS induction, and thus, pol IV concentrations, adding to a growing body of evidence linking ROS and mutational resistance to antibiotics (14,50,60,61). ROS mitigators reduced the number of MuGam foci per cell, indicative of fewer DSBs being formed. ROS accumulation is a major trigger for SOS induction in trimethoprim treated cells and is mediated through RecBCD-dependent resection of ROS-induced DSBs. When a ROS mitigator is including during treatment, the SOS response is not induced even though ssDNA regions are likely to be generated by trimethoprim-induced TLD (5,62,63). Thus, the formation of double-strand breaks is essential for SOS induction in trimethoprim-treated cells. During thymine starvation, ssDNA regions are converted to DSB due to ROS activity (5). Our results indicate that a similar pathway is at play in trimethoprim-treated cells as previously proposed (5).

In ciprofloxacin-treated cells, the deletion of *recB* almost fully inhibited the SOS response. Ciprofloxacin and nalidixic acid both target DNA gyrase (9,11,51). It was previously observed that induction of SOS by the antibiotic nalidixic acid was completely blocked in cells that carried a *recB* mutation and were therefore incapable of processing DSBs through the RecBCD end-resection nuclease complex (36, 37). This implies that SOS induction is also primarily triggered by DSB processing in nalidixic acid-treated cells. Consistent with this result, we showed here that the SOS response in ciprofloxacin-treated cells is *recB*-dependent, consistent with a requirement for DSB processing. Cells lacking *recB* still exhibit very low levels of SOS induction, which could arise from RecA structures assembled on ssDNA regions or by alternative DSB end-resection pathways, for instance *via* a RecJ-dependent pathway proposed previously (64, 65)

Our findings raise the question of whether ssDNA gaps truly represent the major source of SOS induction in *E. coli*. Under our conditions, DSB processing – most often induced by ROS – acts as the major trigger of the SOS response. The results presented here highlight a need that further studies are necessary to fully understand the regulation of the SOS response, in particular the role RecA* structures formed on ssDNA gaps versus DSBs (54,66–68). The observation by Hong *et al*. that ssDNA gaps are converted to DSBs under conditions of thymine starvation (5), highlights ROS-dependent gap-to-break conversion as a potential complicating factor in studies that seek to differentiate events that take place at gaps from those that take place at breaks.

### DSB processing is critical for the formation of pol IV foci

We showed that the processing of ROS-induced DSBs promotes DinB-YPet focus formation. The observations are consistent with a model in which ROS-induced DSBs promote pol IV activity by inducing the SOS response and by generating substrates for pol IV in the form of recombination intermediates.

Few DinB-YPet foci were observed in cells treated with a combination of trimethoprim and DMSO. Based on events that occur during the analogous process of TLD (5, 62), treatment with trimethoprim should induce the formation of ssDNA gaps in the wake of the replisome. In the presence of ROS these would be rapidly converted to DSBs, whereas under ROS mitigated conditions the gaps would persist. The low extent of focus formation observed under trimethoprim-DMSO conditions implies that pol IV rarely acts at these ssDNA gaps.

Following ciprofloxacin treatment, cells exhibited reduced numbers of DinB foci under low ROS conditions. However, ciprofloxacin also induces the formation of end-stabilized DNA-gyrase complexes, which halt DNA synthesis, slowing down cell growth (57, 58). When deleting *recB*, and thus blocking DSB resection at both ROS-induced and ROS-independent DSBs, cells exhibited a very low number of DinB foci, equivalent to numbers present in the absence of damage. Moreover, the colocalization of DinB-YPet with replisomes was substantially increased in the presence of DMSO. It is possible that replisome-proximal DinB-YPet foci, that are insensitive to ROS, reflect pol IV molecules that are recruited to replisomes that have stalled at end-stabilized DNA-gyrase complexes.

### Pol V is not activated by ROS-induced damage

In contrast to the observations made for pol IV, mitigation of ROS produces only a marginal effect on pol V levels in ciprofloxacin-treated cells. Pol V levels barely increase following trimethoprim treatment. Thus unlike pol IV, the repair of ROS-induced DSBs does not directly lead to increased levels of pol V. One possibility is that the mechanisms of SOS induction are different during trimethoprim and ciprofloxacin treatments, with the RecA* structures formed during trimethoprim treatment being insufficient for the up-regulation of pol V. A second and perhaps more likely possibility is that the RecA* structures that trigger LexA cleavage (and thus SOS induction) are different from those that trigger UmuD cleavage (and thus pol V activation). In this scenario, ciprofloxacin treatment may produce both types of RecA* structure, whereas trimethoprim induces only the form competent for SOS induction. In this case, poor cleavage of UmuD would be expected to prevent the accumulation of UmuC due a previously identified system of targeted proteolysis, which limits UmuC accumulation in the absence of UmuD′_2_ (69).

Interestingly, the formation of pol V foci was not affected by adding DMSO to supress DSB formation. This implies that DSBR intermediates are not major substrates for pol V in ciprofloxacin- or trimethoprim-treated cells. In a previous study, we observed that pol V rarely colocalizes with replisomes (47). Together our observations hint at a potential division of labor between pols IV and V, with pol IV often acting at DSBR intermediates and pol V acting at other, as yet unidentified structures, which may include ssDNA gaps or daughter strand gap repair intermediates.

## Materials and Methods

### Strain construction

EAW102 is *E. coli* K-12 MG1655 Δ*recB* and was constructed using λ_RED_ recombination. The kanamycin resistance marker in EAW102 was removed via FLP-FRT recombination (70) using the plasmid pLH29 to obtain kanamycin sensitive HG356.

SSH091, SSH111 and MEC030 (*dinB^+^ lexA^+^ recB^+^* + pUA66-*sulA*-gfp, *dinB^+^ lexA^+^* Δ*recB::FRT* + pUA66-*sulA*-gfp and *recA730 sulA*^-^ + pUA66-*sulA*-gfp) were created by transforming MG1655, EAW102 and EAW287 with pUA66-*sulA*-gfp (49).

RW1286 is *E. coli* MG1655 *umuC-mKate2 dnaX-YPet sulA*^-^::kan^R^ *lexA51*(Def)::Cm^R^ and was made in two steps: first the wild-type *sulA+* gene of EAW282 was replaced with *sulA*^-^::kan by P1 transduction from EAW13 (47), to create EAW282 *sulA*^-^; then *lexA51*(Def) *malB*::Tn*9* was transferred from DE406 (71) into EAW282 *sulA*^-^ by P1 transduction, selecting for chloramphenicol resistance. To confirm the presence of the *lexA*(Def) genotype, colonies were then screened for high levels of RecA expression by Western blotting with anti-RecA antibodies (72).

EAW1144 is *E. coli* K-12 MG1655 *dinB-YPet dnaX-mKate2 sulA*^-^ *lexA51*(Def) Δ*recB* and was constructed in three steps: *sulA*^-^ FRT-Kan-FRT was P1 transduced in EAW643 (KanS) using a P1 lysate grown on EAW13 to obtain the strain EAW1134. The Kan cassette was removed using pLH29(70). Then, *lexA51*(Def) *malB*::Tn*9* was transduced into EAW1134 using a P1 lysate grown on DE406 to obtain the strain EAW1141. Finally, Δ*recB* FRT-KanR-FRT was transduced into EAW1141using P1 lysate grown on EAW102 to obtain EAW1144. All mutations introduced were confirmed by PCR.

The pBAD-*MuGam* vector (pEAW1159) was constructed using a PCR-amplified *muGam* gene fragment (us=GGATATCCATATGGCTAAACCAGCAAAACGTA consisting of a NdeI site and the beginning of the *muGam* gene, and MuGam ds= GCGAATTCTTAAATACCGGCTTCCTGTTCA consisting of an *Eco*RI site and the end of the *muGam* gene) from EAW727 (MG1655 Founder (73) Δe14 with chromosomal *muGam-gfp* in the *att*Tn*7* site). EAW727 was constructed by transducing *muGam-gfp* into Founder Δe14 using a P1 lysate grown on SMR14350 (53). The PCR product was digested with *Nde*I and *Eco*RI and inserted into pBAD *Nde*I which was cut with the same enzymes. pBAD *Nde*I is pBAD/Myc-HisA (Invitrogen) that has been mutated to add a *Nde*I site in place of the original *Nco*I site. All other *Nde*I sites were filled in before the mutagenesis. The resulting plasmid was directly sequenced to confirm presence of wt *muGam* gene

The pBAD-*MuGam-PAmCherry* vector (pEAW1162) was constructed by using two PCR fragments: 1. *Nde*I-MuGam-linker-*Eco*RI generated from pEAW1159 using the following PCR primers: MuGam us=GGATATCCATATGGCTAAACCAGCAAAACGTA consisting of a *Nde*I site and the beginning of the *muGam* gene, and MuGam ds no stop link= GGATATCGAATTCGCCAGAACCAGCAGCGGAGCCAGCGGAAATACCGGCTTCCTGTTC AAATG consisting of an *Eco*RI site, an 11aa linker, and the end of the *muGam* gene without a stop codon. The PCR product was digested with *Nde*I and *Eco*RI. 2. *Eco*RI-PAmCherry-*Hind*III generated from pBAD-*PAmCherry-mCI* (54) using the following PCR primers PAmCherry usEco = GGATATCGAATTCATGGTGAGCAAGGGCGAGGAG consisting of an *Eco*RI site and the beginning of mCherry, and PAmCherry dsHind= GGATATCAAGCTTTTACTTGTACAGCTCGTCCAT consisting of a *Hind*III site and the end of the *mCherry* gene. The PCR product was digested with *Eco*RI and *Hind*III. Both PCR products were ligated to pBAD *Nde*I that had been digested with *Nde*I and *Hind*III. The resulting plasmid was directly sequenced to confirm the presence of *muGam-PAmCherry*.

**Table 1.**
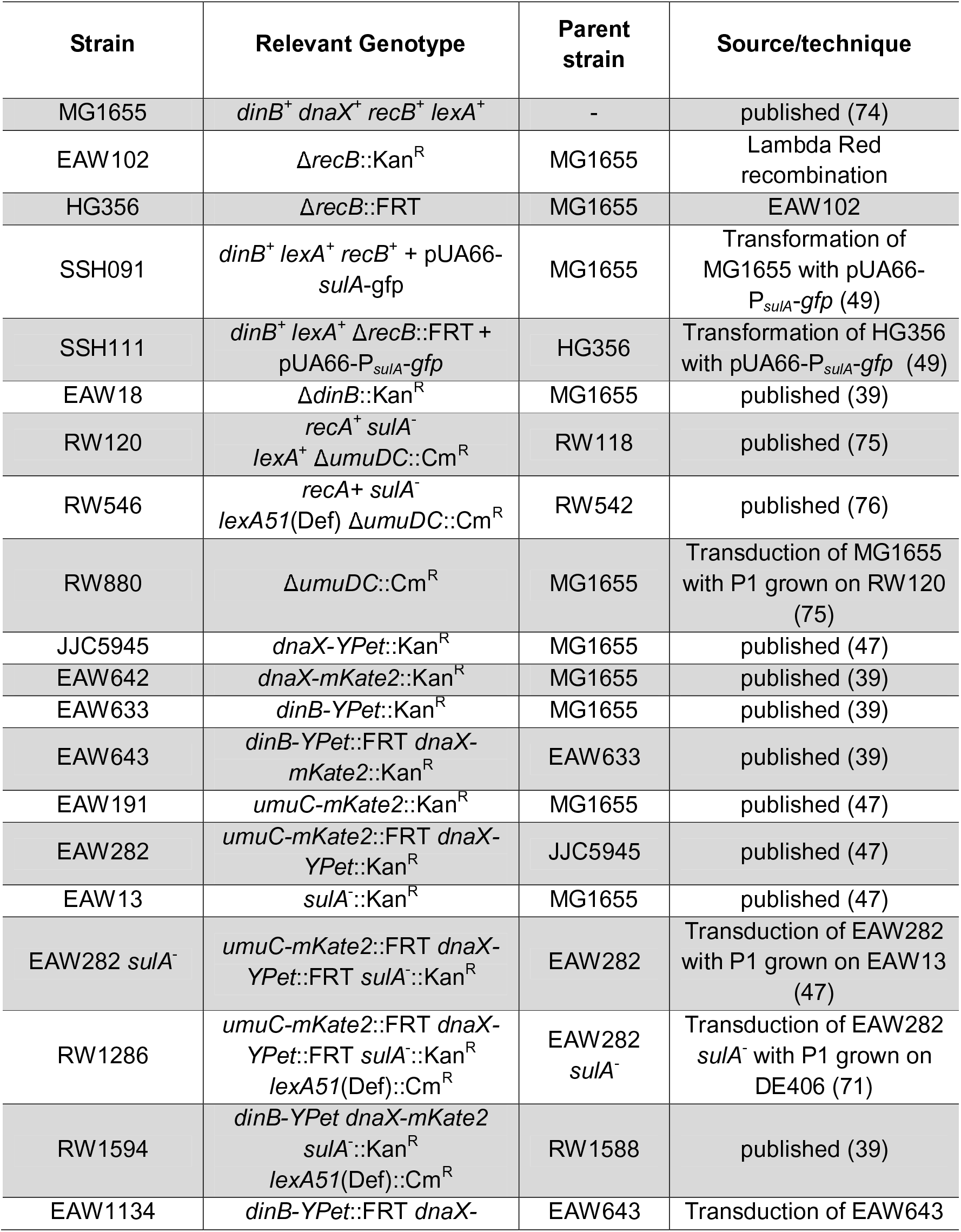
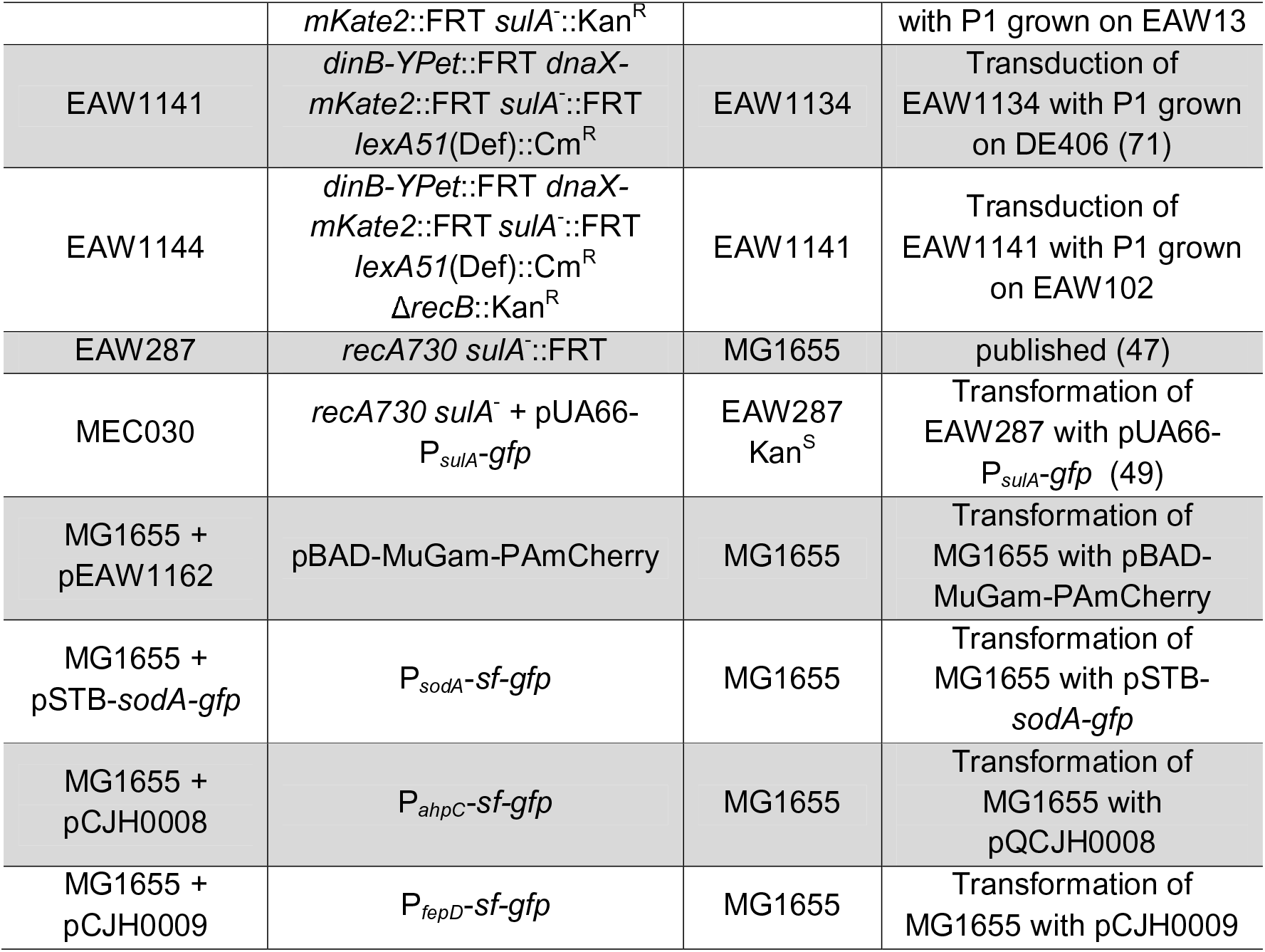
Strains used in this study.

### ROS reporter fusions construction

Three promoters of genes regulated by changes in ROS or iron levels were cloned and fused to the *sf-gfp* gene (45) into a pQBI63 plasmid (Qbiogene). Briefly, upstream regions of *sodA* gene (consisting of the 284 nt intergenic region of *rhaT* and *sodA*) regulated by *soxS* and Fur (4, 41), or *ahpC* gene (−372 to −1 nt of ATG) regulated by OxyR (4,42,43), or *fepD* gene (−170 to −1 nt of ATG) regulated by Fur (44), were amplified and cloned into the pQBI63 plasmid using *Bgl*II/*Nhe*I restriction enzyme to generate respectively pSTB-*sodA-gfp*, pCJH0008 and pCJH0009. All constructions were confirmed by sequencing.

### DNA damaging agent sensitivity assay

Cells were grown in EZ glucose medium overnight at 37°C. The next day, a dilution 1/1000 of each culture was grown in EZ glucose (at 37°C, 150 rpm) until reaching mid log phase (OD_600_ = 0.3). Six aliquots of 300 μL of each culture were transferred in 24 microplates. The first aliquot was used as control of no treatment, 2% DMSO (282 mM, 0.2 x MIC (5)), 30 ng/mL ciprofloxacin, 30 ng/mL ciprofloxacin + 2% DMSO, 1 μg/mL trimethoprim or 1 μg/mL trimethoprim + 2% DMSO were added in the others. Samples of 150 uL were taken at 0 and 60 min; samples at 0 h were taken just before treatment. Each sample was serial diluted in PBS by factor ten down to 10^-6^ and dilutions 10^-1^ to 10^-6^ were spotted on fresh LB plates (Difco brand). Plates were incubated overnight at 37°C in the dark.

### Survival assay following MuGam-PAmCherry expression

To test the effect of MuGam-PAmCherry expression levels on lethality following ciprofloxacin and trimethoprim exposure, seven cells cultures were set up, expressing different levels of MuGam-PAmCherry from a pBAD plasmid. Cells cultures 1-7 (each 1 mL) were grown in EZ glycerol medium in the presence of ampicillin (100 μg/mL) and different L-arabinose concentrations (0, 0.001, 0.003, 0.01, 0.03, 0.1%) and cell culture 8 (1 mL) was grown EZ glucose medium in the presence of ampicillin (100 μg/mL) in overnight at 37°C, 950 rpm. The next day, a 10/1000 dilution of each culture (final volume of 1.5 mL) was grown under the same conditions as over-night growth for 3 h. Each culture was split in three and no drug, 30 ng/mL ciprofloxacin or 1 μg/mL trimethoprim was added. These cultures were grown (at 37°C, 950 rpm) for 2 h. Then, cultures were spin down (5 min; 5,000 g) and cell pellets were resuspended in 0.5 mL corresponding EZ medium; centrifugation and resuspension was carried out three times. Each cell culture was serial diluted in PBS by factor ten down to 10^-5^ and dilutions 10^-1^ to 10^-5^ were spotted on fresh LB plates containing 100 μg/mL ampicillin (Difco brand). Plates were incubated overnight at 37°C in the dark. For each condition, biological triplicates were performed. From these experiments, an L-arabinose concentration of 0.003% was chosen for fluorescence microscopy experiments because this L-arabinose concentration showed no drastic decrease in survival compared to the sample grown in the presence of glucose.

### Plate reader assay

Cells were grown in EZ glucose medium overnight at 37°C. The next day, a dilution 10/1000 of each culture was grown in EZ glucose (at 37°C, 950 rpm) for 3 h. These cultures were diluted to 1/200. Then, 10 μL of these diluted cultures were added to a total volume of 200 μL medium in each well of a 96-well plate. These 200 μL of media contained antibiotic, or hydrogen peroxide, and/or ROS mitigators (final concentration: 5, 10, 20 and 40 ng/mL ± 2% DMSO or ± 0.35 mM BiP; 0.1, 0.3, 1 and 3 μg/mL ± 2% DMSO or ± 0.35 mM BiP; 30, 100, 300 and 500 mM hydrogen peroxide [H_2_O_2_] ± 2% DMSO). For experiments with antibiotics and/or ROS mitigators, antibiotics and/or ROS mitigators were added just before cells were added. For experiments with hydrogen peroxide, hydrogen peroxide was added subsequently after cells were added. For each well, absorbance (OD_600_) is measured every 30 min over 17 h or 18 h. The fluorescence signal was measured at each time point (λ_excitation_ = 470 ± 15 nm, λ_emission_ = 515 ± 20 nm). For cells carrying P*sulA*-gfp, experiments were carried out in 96-well plates from Nalge Nunc International (no. 265301). For cells carrying P*sodA*-sf-gfp, P*ahpC*-sf-gfp or P*fepD*-sf-gfp, experiments were carried out in 96-well plates from Thermo Scientific (no. 165305). The experiments were carried out using the CLARIOstar plate reader (BMG Labtech; settings: orbital reading 4 mm (for 96-well plates from Nalge Nunc International) or 2 mm (for 96-well plates from Thermo Scientific), orbital shaking at 200 rpm, at 37 °C).

Cell cultures were also serial diluted and plated on LB agar plates in order to calculate the number of cells added to each well. To each well, when adding wild-type cells, 10^5^ – 10^6^ cells were added at the beginning of the experiment. For experiments when adding Δ*recB* cells, 10^5^ cells were added at the beginning of the experiment.

### Fluorescence microscopy

For all experiments except for experiments including imaging of MuGam-PAmCherry (Fig. 3), wide-field fluorescence imaging was conducted on an inverted microscope (IX-81, Olympus with a 1.49 NA 100x objective) in an epifluorescence configuration (47). Continuous excitation is provided using semidiode lasers (Sapphire LP, Coherent) of the wavelength 514 nm (150 mW max. output) and 568 nm (200 mW max. output). τ-mKate2 in EAW643 and UmuC-mKate2 in EAW282 were imaged using yellow excitation light (λ = 568 nm) at high intensity (2750 Wcm^-2^), collecting emitted light between 610–680 nm (ET 645/75m filter, Chroma) on a 512 × 512 pixel EM-CCD camera (C9100-13, Hamamatsu). Images of UmuC-mKate2 in RW1286 were recorded at 275 Wcm^-2^. For DinB-YPet imaging of EAW643, we used green excitation (λ = 514 nm) at 160 Wcm^-2^ collecting light emitted between 525–555 nm (ET540/30m filter, Chroma). For DinB-YPet imaging of RW1594, cells were imaged at 51 Wcm^-2^. τ-YPet imaging (EAW282, RW1286) was performed at 51 Wcm^-2^. Cells carrying the SOS reporter plasmid pUA66-*sulA*-gfp (SSH091, SSH111) were imaged at 16 Wcm^-2^.

For experiments including imaging of MuGam-PAmCherry (Fig. 3), imaging was conducted on an inverted microscope (Nikon Eclipse-Ti), equipped with a 1.49 NA 100× objective and a 512 × 512 pixel^2^ Photometrics Evolve CCD camera (Photometrics, Arizona, US). NIS-Elements equipped with JOBS module was used to operate the microscope (Nikon, Japan). Continuous excitation is provided using semidiode lasers of the wavelength 405 nm (OBIS, Coherent, 200 mW max. output) and 568 nm (Sapphire LP, Coherent, 200 mW max. output). MuGam-PAmCherry was imaged by simultaneous illumination with the activation laser 405 nm (1–5 W cm^−2^) and 568 nm readout laser (540 W cm^−2^), a PALM (photoactivation localization microscopy) acquisition protocol, collecting emitted light from 590 nm (ET590LP, Chroma).

Two-color time-lapse movies were recorded to visualize if DinB-YPet foci overlap with τ-mKate2 foci (EAW643). Sets of three images were recorded (bright-field [34 ms exposure], mKate2 fluorescence [100 ms exposure], YPet fluorescence [50 ms exposure]) at an interval of 10 min for 3 h. To measure colocalization between UmuC-mKate2 with the replisome marker τ-YPet (EAW282), we recorded time-lapse movies at the same intervals but different exposures for the replisome marker (bright-field [34 ms exposure], mKate2 fluorescence [100 ms exposure], YPet fluorescence [500 ms exposure]).

Burst acquisitions of DinB-YPet (movies of 300 × 50 ms frames taken every 100 ms light at 514 nm) were collected, subsequently to each burst acquisition, an image of τ-mKate2 (568 nm) was taken (imaging sequence for RW1594). With this imaging sequence, we analysed activity of DinB-YPet at replisomes. RW1286 was imaged similarly; we recorded burst acquisitions of UmuC-mKate2 (568 nm) followed by a snapshot of τ-YPet (514 nm). All images were analysed with ImageJ (77).

The MuGam-PAmCherry imaging acquisition was recorded as a set of two acquisitions, 1. bright-field image (100 ms exposure), 2. PAmCherry fluorescence [simultaneous illumination with the activation laser 405 and 568 nm readout laser for 200 frames each with 100 ms exposure]). This protocol was only executed once for a field-of-view to minimize laser damage. Consequently, before and after antibiotic treatment shows a new set of cells. Images taken after antibiotic addition were recorded following 2 h of antibiotic treatment.

### Flow cell designs

All imaging experiments were carried out in home-built quartz-based flow cells. These flow cells were assembled from a no. 1.5 coverslip (Marienfeld, REF 0102222, for imaging on IX-81, Olympus) or (Marienfeld, REF 0107222, for imaging on Nikon Eclipse-Ti), a quartz top piece (45×20×1 mm) and PE-60 tubing (Instech Laboratories, Inc.). Prior to flow-cell assembly, coverslips were silanized with (3-aminopropyl)triethoxysilane (APTES, from Alfa Aeser). First, coverslips were sonicated for 30 min in a 5M KOH solution to clean and activate the surface. The cleaned coverslips were rinsed thoroughly with MilliQ water and then treated with a 5% (v/v) solution of APTES in MilliQ water. The coverslips were subsequently rinsed with ethanol and sonicated in ethanol for 20 seconds. Afterwards, the coverslips were rinsed with MilliQ water and dried in a jet of N_2_. Silanized slides were stored under vacuum prior to use.

To assemble each flow cell, polyethylene tubing (BTPE-60, Instech Laboratories, Inc.) was glued (BONDiT B-482, Reltek LLC) into two holes that were drilled into a quartz piece. After the glue solidified overnight, double-sided adhesive tape was stuck on two opposite sides of the quartz piece to create a channel. Then, the quartz piece was stuck to an APTES-treated coverslip. The edges were sealed with epoxy glue (5 Minute Epoxy, PARFIX). Each flow cell was stored in a desiccator under mild vacuum while the glue dried. Typical channel dimensions were 45 mm × 5 mm × 0.1 mm (length × width × height).

### Preparation of cell cultures for microscopy

The day before each experiment, for all experiments, an over-night culture was grown from a freezer stock for each cell culture. Cells that did not carry the MuGam-PAmCherry plasmid were grown at 37_⁰_C in EZ rich defined medium (Teknova) that contained 0.2% (w/v) glucose. All strains that have a *Kan*^R^ cassette were grown in the presence of kanamycin (20 μg/mL). Cells that carried the MuGam-PAmCherry plasmid were grown at 37_⁰_C in EZ rich defined medium (Teknova) that contained 0.2% (w/v) glycerol and 0.001% L-arabinose, in the presence of ampicillin (100 μg/mL).

At the day of the experiment, for all imaging experiments excluding imaging of MuGam fusion, cells were grown at 37_⁰_C in EZ rich defined medium (Teknova) that contained 0.2% (w/v) glucose. All strains that have a *Kan*^R^ cassette were grown in the presence of kanamycin (20 μg/mL). Cultures used for imaging under ROS-mitigating conditions were grown in the presence of the particular mitigator used for the experiment (DMSO [2% v/v, 282 mM, 0.2 x MIC (5)] or BiP [0.35 mM, 0.5 x MIC (5)], culture time ∼3 h for *recB*^+^ *lexA*^+^, ∼4 h for Δ*recB lexA*^+^ and ∼6 h for Δ*recB lexA*[Def]). For imaging experiments of the MuGam fusion, cells were grown at 37_⁰_C in EZ rich defined medium (Teknova) that contained 0.2% (w/v) glycerol and 0.001% L-arabinose. All strains were grown in the presence of ampicillin (100 μg/mL). Cultures used for imaging under ROS-mitigating conditions were grown in the presence of DMSO [2% v/v, 282 mM, 0.2 x MIC (5)] for ∼3 h culture time.

### Imaging in flow cells

Cells were loaded into flow cells (*SI Appendix*, Fig. 1*A*), allowed a few minutes to associate with the APTES surface, then loosely associated cells were removed by pulling through fresh medium. The experiment was then initiated by adding either an antibiotic alone or in combination with DMSO to the medium (30 ng/ mL ciprofloxacin, 30 ng/ mL ciprofloxacin with 2% (v/v) DMSO, 1 μg/mL trimethoprim, 1 μg/mL trimethoprim with 2% (v/v) DMSO or 1 μg/mL trimethoprim with 0.35 mM BiP). Throughout the experiment, medium was pulled through the flow cell using a syringe pump, at a rate of 50 μL/min. For each condition, triplicate measurements were recorded.

### Analysis of cell filamentation, concentrations, SOS induction level and number of foci

We selected single cells to obtain information about SOS induction, DinB and UmuC levels upon UV irradiation (>100 cells for every time point). MicrobeTracker 0.937 (78), a MATLAB script, was used to create cell outlines as regions of interest (ROI). We manually curated cell outlines designated by MicrobeTracker at t = 0 min (time point of antibiotic addition) and at 30 min time intervals until 180 min. By obtaining cell outlines manually, we ensure accuracy and purely select non-overlapping, in-focus cells for analysis. These ROI were imported in ImageJ 1.50i. The cell outlines were then used to measure mean cell intensities, cell lengths and the number of foci per cell. Parameters describing foci (number, positions and intensities) were obtained using a Peak Fitter plug-in, described previously (39, 47). Prior to determining DinB-YPet foci UmuC-mKate2 per cell from burst acquisition movies in *lexA*(Def), average projections in time were curated from frame 1 to 101 (10 x 100 ms = 1 s). Prior to determining MuGam-PAmCherry foci per cell from burst acquisition movies, maximum projections in time were curated over the entire movie, capturing all binding events of MuGam-PAmCherry.

Using information of mean cell brightness derived from DinB-YPet expressing cells, we also calculated DinB-YPet concentrations of cells grown in the absence or presence of antibiotic. In a previous study (39), we calculated the DinB-YPet concentration which correlates with a certain mean cell brightness (in the absence of ciprofloxacin: 6 ± 1 nm [SE]; 180 min after ciprofloxacin treatment: 34 ± 3 nM [SE]). We utilized these values to calculate the DinB-YPet concentration for ciprofloxacin ± DMSO or trimethoprim ± DMSO treated cells.

### Analysis of colocalization events

Foci were classed as colocalized if their centroid positions (determined using our peak fitter tool) fell within 2.18 px (218 nm) of each other. When treating with ciprofloxacin, we determined that for DinB-YPet–τ-mKate2 localization the background of DinB foci expected to colocalize with replisomes purely by chance is ∼4% at 180 min. This was calculated by taking the area of each cell occupied by replisome foci (including the colocalization search radius) and dividing by the total area of the cell. The value of 4% corresponds to the mean of measurements made over 121 cells. Since the foci density of replisomes stays fairly constant following ciprofloxacin treatment, the chance colocalization of DinB-YPet foci with τ-mKate2 is ∼4% during the experiment (39). Chance colocalization of τ-mKate2 with DinB-YPet is however not constant over time because most cells contain no pol IV foci in the absence of any DNA damage. Chance colocalization is close to zero at 0 min; at 60 min, chance colocalization is ∼5%; at 120 min, chance colocalization is ∼3%. Moreover, chance colocalization of τ-mKate2 with DinB-YPet is overall reduced under ROS-mitigating conditions due to a reduced number of foci per cell (chance colocalization close to zero at 0 min; at 120 min, ∼2%). Chance colocalization of τ-mKate2 with DinB-YPet in trimethoprim-treated cells amounts to ∼1% from 60-90 min (close to zero before 60 min). Under ROS-mitigating conditions, chance colocalization is always close to zero because the number of pol IV foci per cell does not increase post treatment as well as cell size (Fig. 1).

The chance colocalization of UmuC-mKate2 with τ-YPet is similar to the chance colocalization of DinB-YPet with τ-mKate2 (chance colocalization: ∼4%). The expected colocalization of τ-YPet with UmuC-mKate2 by background is close to zero until 90 min. UmuC-mKate2 is neither upregulated nor released from the membrane (*SI Appendix*, Fig. 13*A*). Chance colocalization is ∼3% at 180 min after ciprofloxacin treatment and ∼2% after the combinational treatment of ciprofloxacin/DMSO.

### Western blotting

Overnight *E. coli* LB cultures of RW120/pRW154 and RW546/pRW154 (75) were diluted 1 to 100 in fresh LB with appropriate antibiotics and grown to mid-log (∼OD 0.5, ∼3 hrs). Aliquots were then taken for the untreated samples. Either ciprofloxacin (30 ng/mL) or trimethoprim (1 µg/mL) was added to the remaining culture and incubated with or without the addition of 2% DMSO. Samples were taken at 1, 2 and 3 hours. Whole cell extracts were made by centrifuging 1.5 mL of culture and adding 90 µl of sterile deionized water and 30µL of NuPAGE LDS sample buffer (4X) (Novex, Life Technologies) to the cell pellet. Five cycles of freeze/thaw on dry ice and in a 37_⁰_C water bath were performed to lyse the cells. Extracts were boiled for 5 minutes prior to loading. Samples were run on NuPAGE 4-12% Bis-Tris gels (Novex Life Technologies) and transferred to Invitrolon PVDF (0.45 µm pore size) membranes (Novex Life Technologies). Membranes were incubated with anti-UmuD antibodies (1:5,000 dilution) at room temperature overnight. Then the membranes were incubated with goat anti-rabbit IgG (H+L) alkaline phosphatase conjugate (1:10,000 dilution) (BIO-RAD). Subsequently, the membranes were treated with the CDP-Star substrate (Tropix). Membranes were then exposed to BioMax XAR film (Carestream) to visualize UmuD protein bands.

## Acknowledgements

We thank Amy E. McGrath, Jonathan Williams, Martina L. Sanderson-Smith for assistance with plate reader assays. MMC was supported by grant GM32335 from the National Institute of General Medical Sciences USA. RW was supported by funds from the National Institutes of Health, National Institute of Child Health and Human Development Intramural Research Program. AvO was supported by a Laureate Fellowship FL140100027 from the Australian Research Council. AR was supported by was supported by Project Grant APP1165135 from the National Health and Medical Research Council and funds from the Faculty of Science, Medicine and Health and the Illawarra Health and Medical Research Institute.

## Supplementary Information Text

**<u>Sequence of pBAD-MuGam-PAmCherry (pEAW1162)</u>.** AAGAAACCAATTGTCCATATTGCATCAGACATTGCCGTCACTGCGTCTTTTACTGGCTCT TCTCGCTAACCAAACCGGTAACCCCGCTTATTAAAAGCATTCTGTAACAAAGCGGGACC AAAGCCATGACAAAAACGCGTAACAAAAGTGTCTATAATCACGGCAGAAAAGTCCACAT TGATTATTTGCACGGCGTCACACTTTGCTATGCCATAGCATTTTTATCCATAAGATTAGC GGATCCTACCTGACGCTTTTTATCGCAACTCTCTACTGTTTCTCCATACCCGTTTTTTGG GCTAACAGGAGGAATTAACATATGGCTAAACCAGCAAAACGTATCAAGAGTGCCGCAG CGGCTTATGTGCCACAAAACCGCGATGCGGTGATTACCGATATTAAACGCATCGGGGA TTTACAGCGCGAAGCATCACGTCTGGAAACGGAAATGAATGATGCCATCGCGGAAATTA CGGAGAAATTTGCGGCCCGGATTGCACCGATTAAAACCGATATTGAAACCCTTTCAAAA GGCGTTCAGGGATGGTGTGAAGCGAACCGCGACGAACTGACGAACGGCGGCAAAGTG AAGACGGCGAATCTTGTCACCGGTGATGTATCGTGGCGGGTCCGTCCACCATCAGTAA GTATTCGTGGTATGGATGCAGTGATGGAAACGCTGGAGCGTCTTGGCCTGCAACGCTT TATTCGCACGAAGCAGGAAATCAACAAGGAAGCGATTTTACTGGAACCGAAAGCGGTC GCAGGCGTTGCCGGAATTACAGTTAAATCAGGCATTGAGGATTTTTCTATTATTCCATTT GAACAGGAAGCCGGTATTTCCGCTGGCTCCGCTGCTGGTTCTGGCGAATTCATGGTGA <u>GCAAGGGCGAGGAGGATAACATGGCCATCATTAAGGAGTTCATGCGCTTCAAGGTGCA</u> <u>CATGGAGGGGTCCGTGAACGGCCACGTGTTCGAGATCGAGGGCGAGGGCGAGGGCC</u> <u>GCCCCTACGAGGGCACCCAGACCGCCAAGCTGAAGGTGACCAAGGGTGGCCCCCTGC</u> <u>CCTTCACCTGGGACATCCTGTCCCCTCAATTCATGTACGGCTCCAATGCCTACGTGAAG</u> <u>CACCCCGCCGACATCCCCGACTACTTTAAGCTGTCCTTCCCCGAGGGCTTCAAGTGGG</u> <u>AGCGCGTGATGAAATTCGAGGACGGCGGCGTGGTGACCGTGACCCAGGACTCCTCCC</u> <u>TGCAGGACGGTGAGTTCATCTACAAGGTGAAGCTGCGCGGCACCAACTTCCCCTCCGA</u> <u>CGGCCCCGTAATGCAGAAGAAGACCATGGGCTGGGAGGCCCTCTCCGAGCGGATGTA</u> <u>CCCCGAGGACGGCGCCCTGAAGGGCGAGGTCAAGCCGAGAGTGAAGCTGAAGGACG</u> <u>GCGGCCACTACGACGCTGAGGTCAAGACCACCTACAAGGCCAAGAAGCCCGTGCAGC</u> <u>TGCCCGGCGCCTACAACGTCAACCGCAAGTTGGACATCACCTCACACAACGAGGACTA</u> <u>CACCATCGTGGAACAGTACGAACGTGCCGAGGGCCGCCACTCCACCGGCGGCATGGA</u> CGAGCTGTACAAGTAAAAGCTTGGGCCCGAACAAAAACTCATCTCAGAAGAGGATCTG AATAGCGCCGTCGACCATCATCATCATCATCATTGAGTTTAAACGGTCTCCAGCTTGGC TGTTTTGGCGGATGAGAGAAGATTTTCAGCCTGATACAGATTAAATCAGAACGCAGAAG CGGTCTGATAAAACAGAATTTGCCTGGCGGCAGTAGCGCGGTGGTCCCACCTGACCCC ATGCCGAACTCAGAAGTGAAACGCCGTAGCGCCGATGGTAGTGTGGGGTCTCCCCATG CGAGAGTAGGGAACTGCCAGGCATCAAATAAAACGAAAGGCTCAGTCGAAAGACTGGG CCTTTCGTTTTATCTGTTGTTTGTCGGTGAACGCTCTCCTGAGTAGGACAAATCCGCCG GGAGCGGATTTGAACGTTGCGAAGCAACGGCCCGGAGGGTGGCGGGCAGGACGCCC GCCATAAACTGCCAGGCATCAAATTAAGCAGAAGGCCATCCTGACGGATGGCCTTTTTG CGTTTCTACAAACTCTTTTGTTTATTTTTCTAAATACATTCAAATATGTATCCGCTCATGA GACAATAACCCTGATAAATGCTTCAATAATATTGAAAAAGGAAGAGTATGAGTATTCAAC ATTTCCGTGTCGCCCTTATTCCCTTTTTTGCGGCATTTTGCCTTCCTGTTTTTGCTCACC CAGAAACGCTGGTGAAAGTAAAAGATGCTGAAGATCAGTTGGGTGCACGAGTGGGTTA CATCGAACTGGATCTCAACAGCGGTAAGATCCTTGAGAGTTTTCGCCCCGAAGAACGTT TTCCAATGATGAGCACTTTTAAAGTTCTGCTATGTGGCGCGGTATTATCCCGTGTTGAC GCCGGGCAAGAGCAACTCGGTCGCCGCATACACTATTCTCAGAATGACTTGGTTGAGT ACTCACCAGTCACAGAAAAGCATCTTACGGATGGCATGACAGTAAGAGAATTATGCAGT GCTGCCATAACCATGAGTGATAACACTGCGGCCAACTTACTTCTGACAACGATCGGAG GACCGAAGGAGCTAACCGCTTTTTTGCACAACATGGGGGATCATGTAACTCGCCTTGAT CGTTGGGAACCGGAGCTGAATGAAGCCATACCAAACGACGAGCGTGACACCACGATG CCTGTAGCAATGGCAACAACGTTGCGCAAACTATTAACTGGCGAACTACTTACTCTAGC TTCCCGGCAACAATTAATAGACTGGATGGAGGCGGATAAAGTTGCAGGACCACTTCTG CGCTCGGCCCTTCCGGCTGGCTGGTTTATTGCTGATAAATCTGGAGCCGGTGAGCGTG GGTCTCGCGGTATCATTGCAGCACTGGGGCCAGATGGTAAGCCCTCCCGTATCGTAGT TATCTACACGACGGGGAGTCAGGCAACTATGGATGAACGAAATAGACAGATCGCTGAG ATAGGTGCCTCACTGATTAAGCATTGGTAACTGTCAGACCAAGTTTACTCATATATACTT TAGATTGATTTAAAACTTCATTTTTAATTTAAAAGGATCTAGGTGAAGATCCTTTTTGATA ATCTCATGACCAAAATCCCTTAACGTGAGTTTTCGTTCCACTGAGCGTCAGACCCCGTA GAAAAGATCAAAGGATCTTCTTGAGATCCTTTTTTTCTGCGCGTAATCTGCTGCTTGCAA ACAAAAAAACCACCGCTACCAGCGGTGGTTTGTTTGCCGGATCAAGAGCTACCAACTCT TTTTCCGAAGGTAACTGGCTTCAGCAGAGCGCAGATACCAAATACTGTCCTTCTAGTGT AGCCGTAGTTAGGCCACCACTTCAAGAACTCTGTAGCACCGCCTACATACCTCGCTCTG CTAATCCTGTTACCAGTGGCTGCTGCCAGTGGCGATAAGTCGTGTCTTACCGGGTTGG ACTCAAGACGATAGTTACCGGATAAGGCGCAGCGGTCGGGCTGAACGGGGGGTTCGT GCACACAGCCCAGCTTGGAGCGAACGACCTACACCGAACTGAGATACCTACAGCGTGA GCTATGAGAAAGCGCCACGCTTCCCGAAGGGAGAAAGGCGGACAGGTATCCGGTAAG CGGCAGGGTCGGAACAGGAGAGCGCACGAGGGAGCTTCCAGGGGGAAACGCCTGGT ATCTTTATAGTCCTGTCGGGTTTCGCCACCTCTGACTTGAGCGTCGATTTTTGTGATGCT CGTCAGGGGGGCGGAGCCTATGGAAAAACGCCAGCAACGCGGCCTTTTTACGGTTCC TGGCCTTTTGCTGGCCTTTTGCTCACATGTTCTTTCCTGCGTTATCCCCTGATTCTGTGG ATAACCGTATTACCGCCTTTGAGTGAGCTGATACCGCTCGCCGCAGCCGAACGACCGA GCGCAGCGAGTCAGTGAGCGAGGAAGCGGAAGAGCGCCTGATGCGGTATTTTCTCCT TACGCATCTGTGCGGTATTTCACACCGCATAtaTGGTGCACTCTCAGTACAATCTGCTCT GATGCCGCATAGTTAAGCCAGTATACACTCCGCTATCGCTACGTGACTGGGTCATGGC TGCGCCCCGACACCCGCCAACACCCGCTGACGCGCCCTGACGGGCTTGTCTGCTCCC GGCATCCGCTTACAGACAAGCTGTGACCGTCTCCGGGAGCTGCATGTGTCAGAGGTTT TCACCGTCATCACCGAAACGCGCGAGGCAGCAGATCAATTCGCGCGCGAAGGCGAAG CGGCATGCATAATGTGCCTGTCAAATGGACGAAGCAGGGATTCTGCAAACCCTATGCT ACTCCGTCAAGCCGTCAATTGTCTGATTCGTTACCAATTATGACAACTTGACGGCTACAT CATTCACTTTTTCTTCACAACCGGCACGGAACTCGCTCGGGCTGGCCCCGGTGCATTTT TTAAATACCCGCGAGAAATAGAGTTGATCGTCAAAACCAACATTGCGACCGACGGTGG CGATAGGCATCCGGGTGGTGCTCAAAAGCAGCTTCGCCTGGCTGATACGTTGGTCCTC GCGCCAGCTTAAGACGCTAATCCCTAACTGCTGGCGGAAAAGATGTGACAGACGCGAC GGCGACAAGCAAACATGCTGTGCGACGCTGGCGATATCAAAATTGCTGTCTGCCAGGT GATCGCTGATGTACTGACAAGCCTCGCGTACCCGATTATCCATCGGTGGATGGAGCGA CTCGTTAATCGCTTCCATGCGCCGCAGTAACAATTGCTCAAGCAGATTTATCGCCAGCA GCTCCGAATAGCGCCCTTCCCCTTGCCCGGCGTTAATGATTTGCCCAAACAGGTCGCT GAAATGCGGCTGGTGCGCTTCATCCGGGCGAAAGAACCCCGTATTGGCAAATATTGAC GGCCAGTTAAGCCATTCATGCCAGTAGGCGCGCGGACGAAAGTAAACCCACTGGTGAT ACCATTCGCGAGCCTCCGGATGACGACCGTAGTGATGAATCTCTCCTGGCGGGAACAG CAAAATATCACCCGGTCGGCAAACAAATTCTCGTCCCTGATTTTTCACCACCCCCTGAC CGCGAATGGTGAGATTGAGAATATAACCTTTCATTCCCAGCGGTCGGTCGATAAAAAAA TCGAGATAACCGTTGGCCTCAATCGGCGTTAAACCCGCCACCAGATGGGCATTAAACG AGTATCCCGGCAGCAGGGGATCATTTTGCGCTTCAGCCATACTTTTCATACTCCCGCCA TTCAGAG

---

**Fig. S1.**
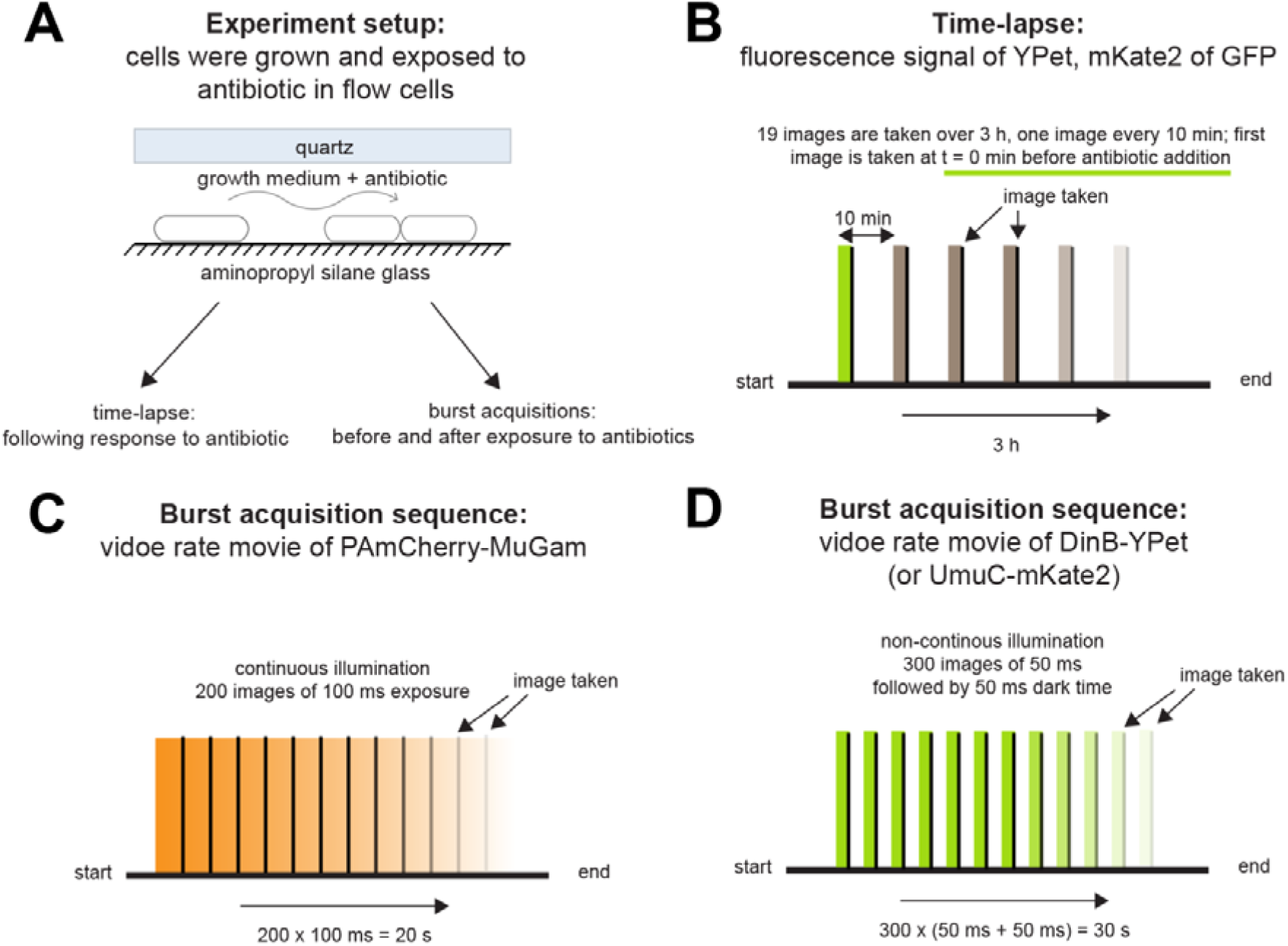
Experimental design. (*A*) Experimental setup. Cells are loaded in a flow-cell and immobilized on a positively charged aminopropyl silane glass surface. Cells were imaged before and after antibiotic exposure ± ROS mitigator. Time-lapse movies were recorded to follow the cellular response. Burst acquisitions were recorded to follow the dynamic behavior of fluorescent protein fusion constructs in cells. (*B*) Time-lapse movies were recorded over 3 h following the cellular response to antibiotic exposure. An image was taken every 10 min. At t = 0 min, the first image was taken and subsequently antibiotic-containing media was flowed into the flow cell. A total number of 19 frames were recorded. (*C*) Burst acquisition videos were recorded at specific time-points before or after antibiotic addition. Movies of MuGam-PAmCherry were recorded using continuous excitation, containing 200 frames at 100 ms exposure. (*D*) Burst acquisition movies of DinB-YPet or UmuC-mKate2 were recorded using non-continuous excitation, containing 300 frames at 50 ms exposure followed by 50 ms dark time.

**Fig. S2.**
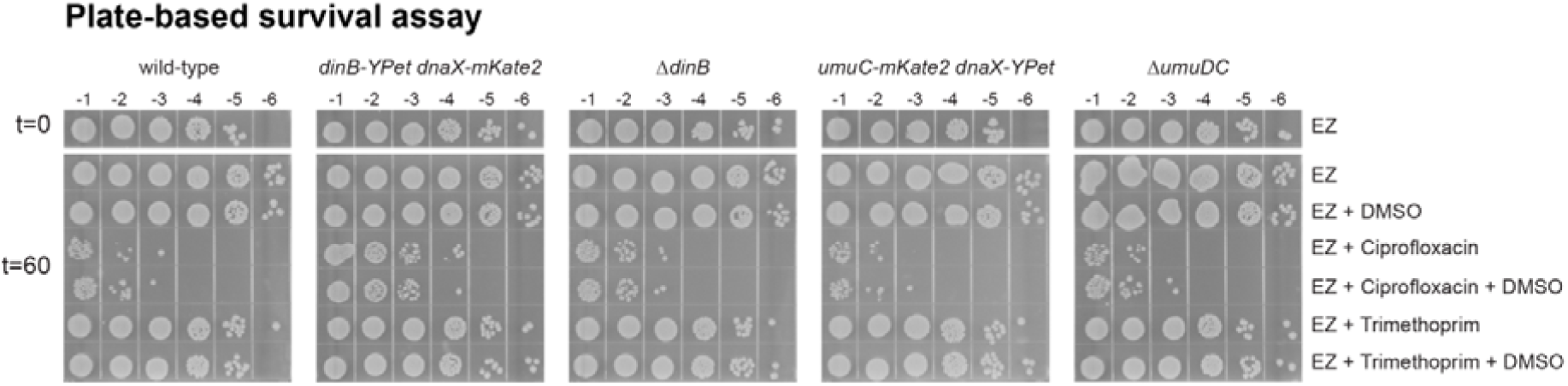
Survival of strains to ciprofloxacin and trimethoprim in EZ medium. Survival assays using ciprofloxacin or trimethoprim normal or ROS-mitigating condition (+ DMSO). Cell cultures (MG1655 [wild-type], *dinB-YPet dnaX-mKate2*, Δ*dinB*, *umuC-mKate2 dnaX-YPet* and Δ*umuDC*) were grown in EZ glucose medium to exponential growth phase (OD_600_ = 0.2-0.3). Then, culture were split in 6 before, one sample was used as control, 2% DMSO, 30 ng/mL ciprofloxacin, 30 ng/mL ciprofloxacin + DMSO, 1 μg/mL trimethoprim or or 1 μg/mL trimethoprim + 2% DMSO were added in the others and grown for 60 min. Before the treatment and after 60 min samples were taken and serial diluted by factor ten down to 10^-6^. Dilutions 10^-1^ to 10^-6^ of each culture were spotted on fresh LB plates, incubated in the dark overnight at 37°C before the image were captured. Images selected are resentative of a biological triplicate. Cells constructs used in this study (*dinB-YPet dnaX-mKate2* and *umuC-mKate2 dnaX-YPet*) exhibit a similar phenotype to MG1655.

**Fig. S3.**
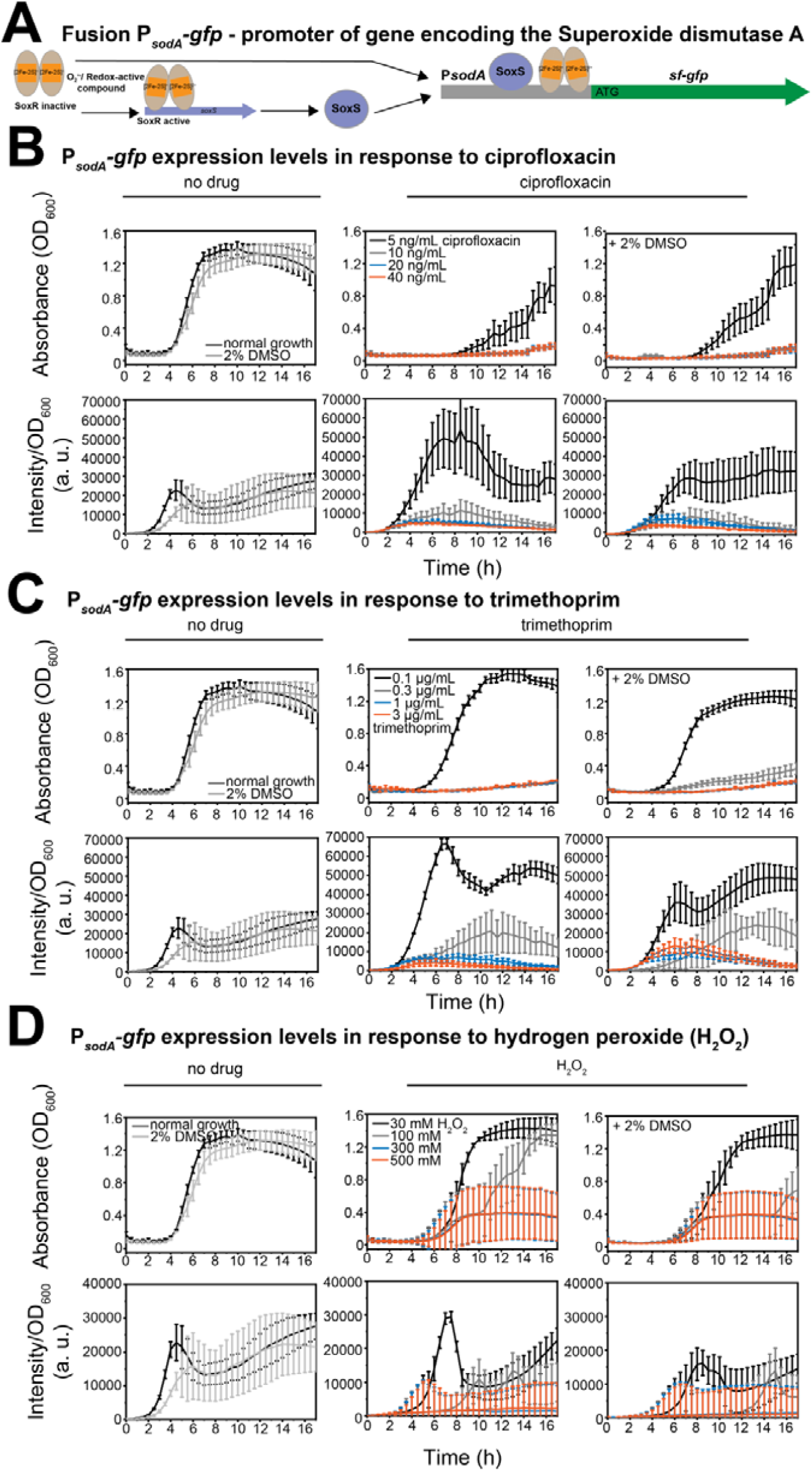
P*_sodA_-gfp* expression levels wild-type cells. For each strain, 10^4^ – 10^6^ cells were added to each well at the beginning of the experiment. Measurements of absorbance (OD_600_) and fluorescence intensity (a.u.) were carried out every 30 min over 17 h. For (A)-(C): upper row shows absorbance (OD_600_) and bottom row illustrates intensity values/ OD_600_, consistent with expression levels. Error bars represent standard error of the mean over three independent biological replicates. (*A*) *sodA* is regulated by SoxRS. Superoxides oxidize the Fe-S clusters of the SoxR transcription factor, promoting transcription of *soxS* and *sodA*. Then, SoxS also acts as a transcription factor for *sodA*. For cells carrying P*_sodA_-gfp*, superoxides then trigger the expression of GFP from the *sodA* promotor. (*B*) Comparison of normal growth condition with ciprofloxacin treatment ± ROS mitigator for wild-type cells. First column: normal growth conditions (wild-type: dark grey; Δ*recB*: orange) or + 2% DMSO (wild-type: grey); second column: ciprofloxacin treatment of wild-type cells (5 ng/mL: black; 10 ng/mL: grey; 20 ng/mL: blue; 40 ng/mL: orange); third column: ciprofloxacin + 2% DMSO treatment of wild-type cells (same color coding as second column). (*C*) Comparison of normal growth condition with trimethoprim treatment ± ROS mitigator for wild-type cells. First column: as (A) first column; second column: trimethoprim treatment of wild-type cells (0.1 μg/mL: black; 0.3 μg/mL: grey; 1 μg/mL: blue; 3 μg/mL: orange); third column: trimethoprim + 2% DMSO treatment of wild-type cells (same color coding as second column). (*D*) Comparison of normal growth condition with hydrogen peroxide (H_2_O_2_) treatment ± ROS mitigator for wild-type cells. First column: as (A) first column; second column: H_2_O_2_ treatment of wild-type cells (30 mM: black; 100 mM: grey; 300 mM: blue; 500 mM: orange); third column: H_2_O_2_ + 2% DMSO treatment of wild-type cells (same color coding as second column).

**Fig. S4.**
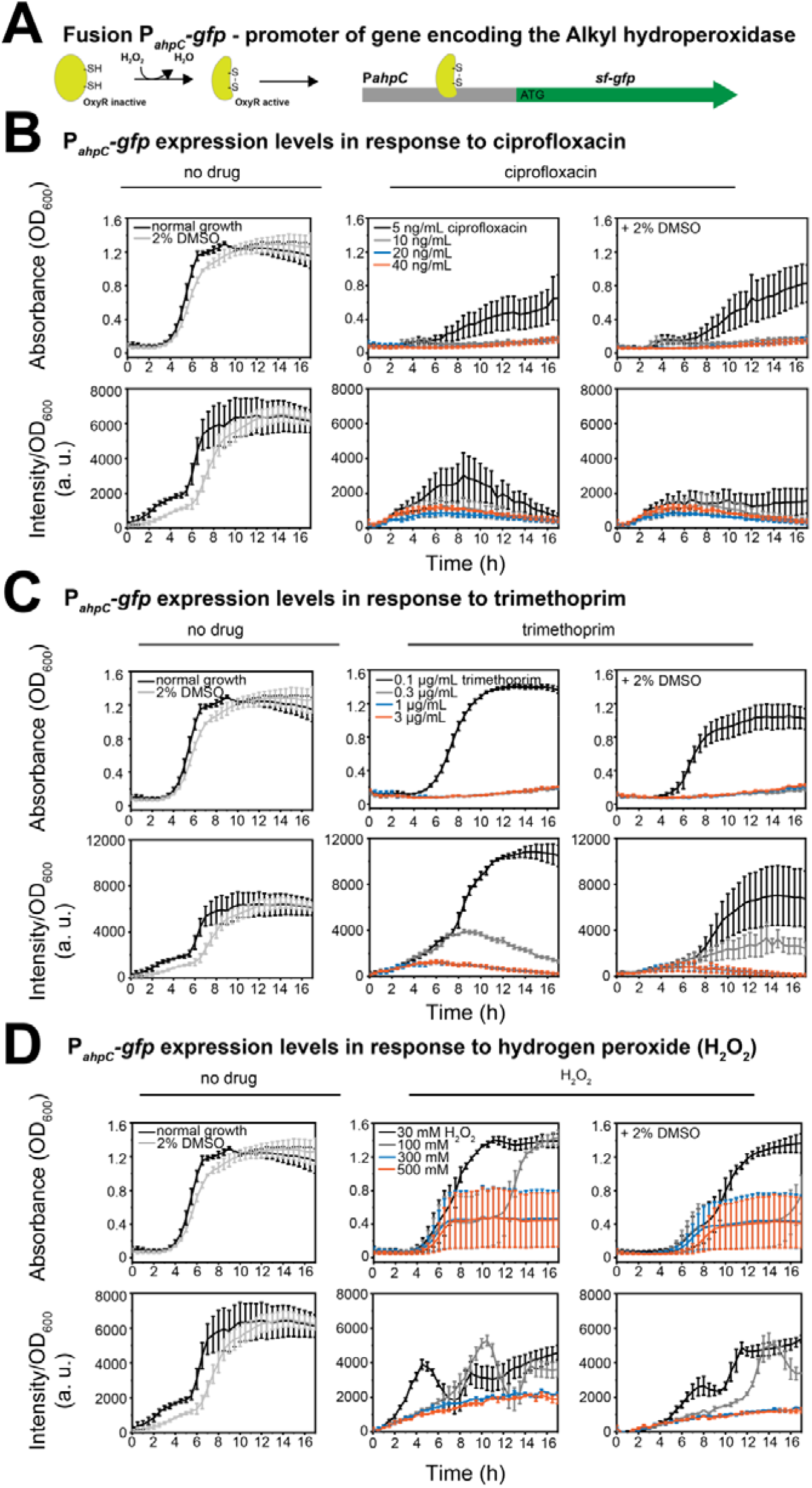
P*_ahpC_-gfp* expression levels wild-type cells. For each strain, 10^4^ – 10^6^ cells were added to each well at the beginning of the experiment. Measurements of absorbance (OD_600_) and fluorescence intensity (a.u.) were carried out every 30 min over 17 h. For (A)-(C): upper row shows absorbance (OD_600_) and bottom row illustrates intensity values/ OD_600_, consistent with expression levels. Error bars represent standard error of the mean over three independent biological replicates. (*A*) *ahcP* is transcriptionally regulated by OxyR. Oxidation of OxyR cysteines induces transcription and expression of *ahcPC*. For cells carrying P*_ahcP_-gfp*, oxidative stress triggers the expression of GFP from the *ahcP* promotor. (*B*) Comparison of normal growth condition with ciprofloxacin treatment ± ROS mitigator for wild-type cells. First column: normal growth conditions (wild-type: dark grey; Δ*recB*: orange) or + 2% DMSO (wild-type: grey); second column: ciprofloxacin treatment of wild-type cells (5 ng/mL: black; 10 ng/mL: grey; 20 ng/mL: blue; 40 ng/mL: orange); third column: ciprofloxacin + 2% DMSO treatment of wild-type cells (same color coding as second column). (*C*) Comparison of normal growth condition with trimethoprim treatment ± ROS mitigator for wild-type cells. First column: as (A) first column; second column: trimethoprim treatment of wild-type cells (0.1 μg/mL: black; 0.3 μg/mL: grey; 1 μg/mL: blue; 3 μg/mL: orange); third column: trimethoprim + 2% DMSO treatment of wild-type cells (same color coding as second column). (*D*) Comparison of normal growth condition with hydrogen peroxide (H_2_O_2_) treatment ± ROS mitigator for wild-type cells. First column: as (A) first column; second column: H_2_O_2_ treatment of wild-type cells (30 mM: black; 100 mM: grey; 300 mM: blue; 500 mM: orange); third column: H_2_O_2_ + 2% DMSO treatment of wild-type cells (same color coding as second column).

**Fig. S5.**
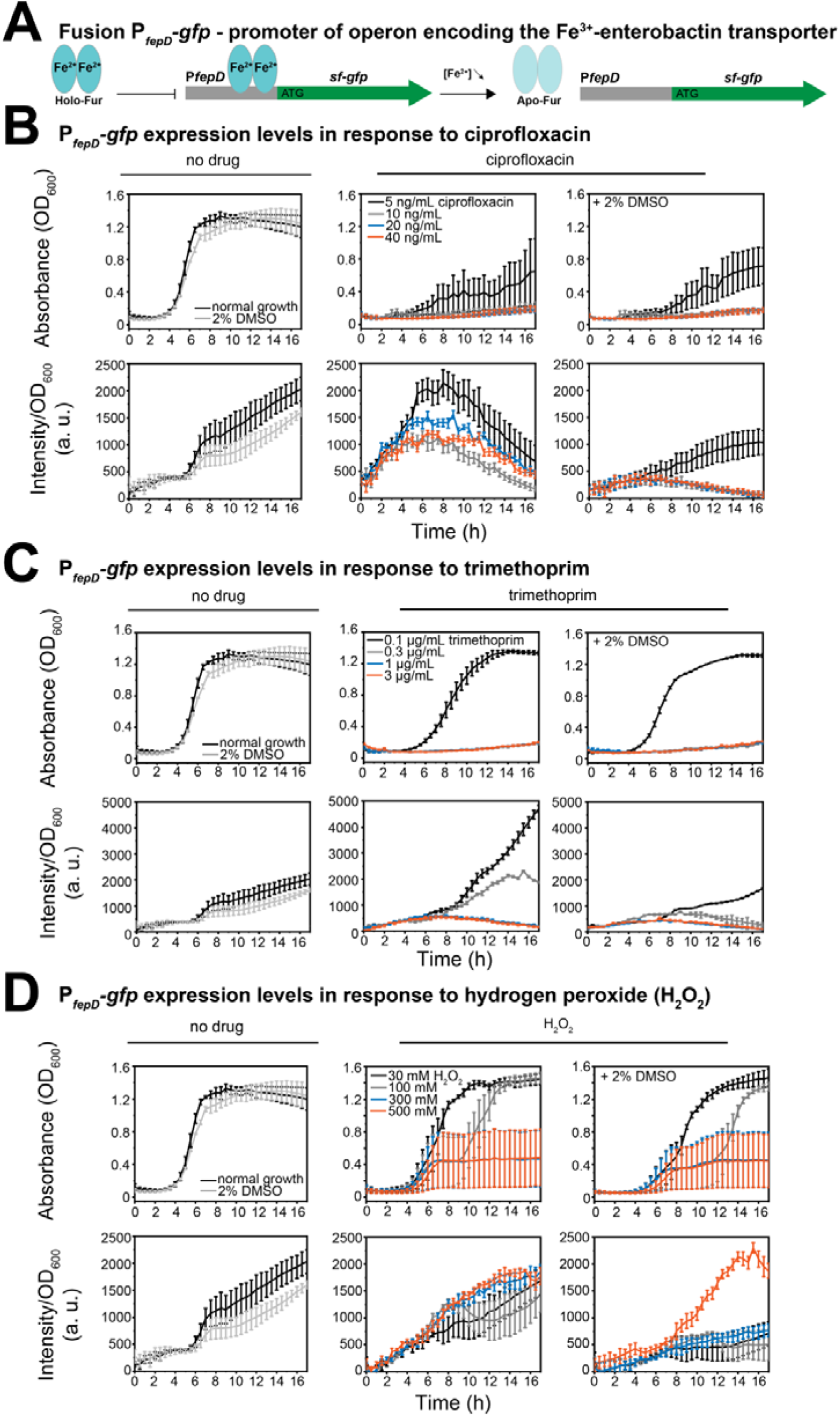
P*_fepD_-gfp* expression levels wild-type cells. For each strain, 10^4^ – 10^6^ cells were added to each well at the beginning of the experiment. Measurements of absorbance (OD_600_) and fluorescence intensity (a.u.) were carried out every 30 min over 17 h. For (A)-(C): upper row shows absorbance (OD_600_) and bottom row illustrates intensity values/ OD_600_, consistent with expression levels. Error bars represent standard error of the mean over three independent biological replicates. (*A*) *fepD* is regulated by Fur. Under high iron conditions, transcriptional repressor Fur inhibits of *fepD* transcription. Under low iron conditions, in the presence of oxidative damage, Fur is de-repressed and *fepD* is transcribed. For cells carrying P*_fepD_-gfp*, oxidative stress triggers the expression of GFP from the *fepD* promotor. (*B*) Comparison of normal growth condition with ciprofloxacin treatment ± ROS mitigator for wild-type cells. First column: normal growth conditions (wild-type: dark grey; Δ*recB*: orange) or + 2% DMSO (wild-type: grey); second column: ciprofloxacin treatment of wild-type cells (5 ng/mL: black; 10 ng/mL: grey; 20 ng/mL: blue; 40 ng/mL: orange); third column: ciprofloxacin + 2% DMSO treatment of wild-type cells (same color coding as second column). (*C*) Comparison of normal growth condition with trimethoprim treatment ± ROS mitigator for wild-type cells. First column: as (A) first column; second column: trimethoprim treatment of wild-type cells (0.1 μg/mL: black; 0.3 μg/mL: grey; 1 μg/mL: blue; 3 μg/mL: orange); third column: trimethoprim + 2% DMSO treatment of wild-type cells (same color coding as second column). (*D*) Comparison of normal growth condition with hydrogen peroxide (H_2_O_2_) treatment ± ROS mitigator for wild-type cells. First column: as (A) first column; second column: H_2_O_2_ treatment of wild-type cells (30 mM: black; 100 mM: grey; 300 mM: blue; 500 mM: orange); third column: H_2_O_2_ + 2% DMSO treatment of wild-type cells (same color coding as second column).

**Fig. S6.**
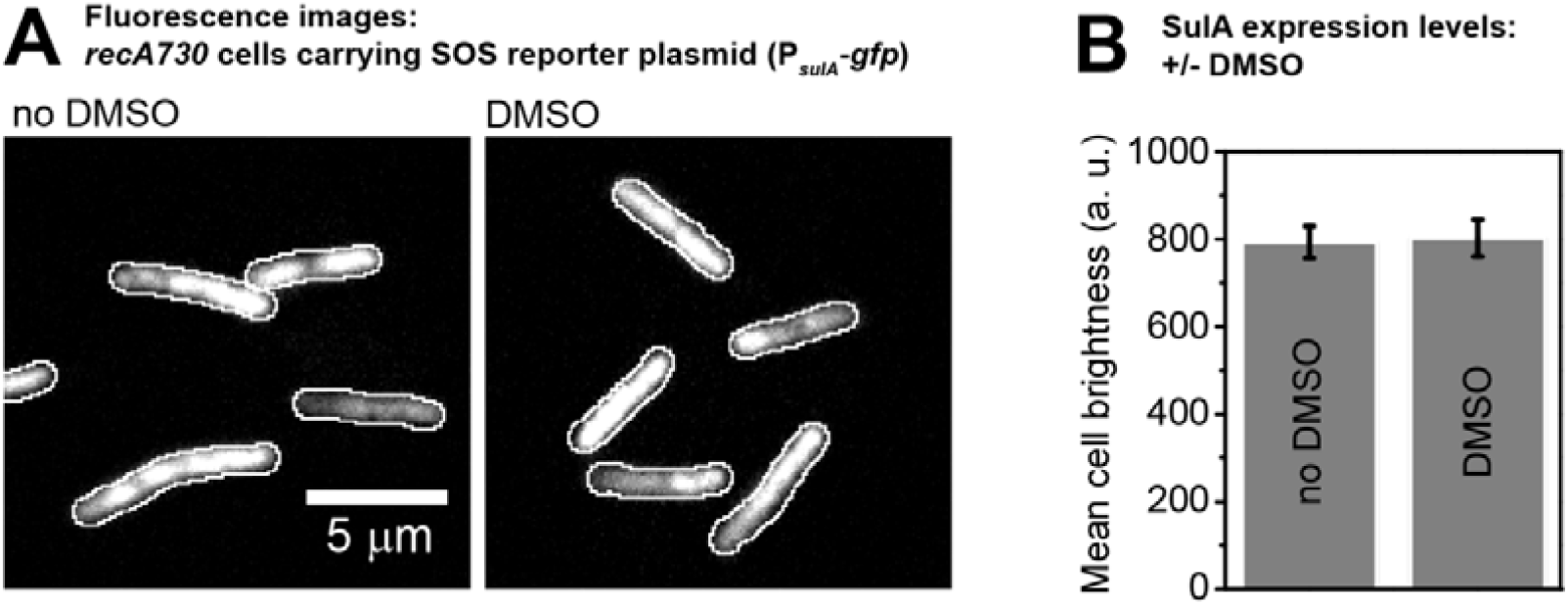
DMSO has no effect on GFP fluorescence *in vivo*. (*A*) Fluorescence images of *recA730* cells carrying the SOS reporter plasmid (P*_sulA_-gfp*) in the absence of DMSO (left) and in the presence of DMSO (right). Scale bar represents 5 µm. (*B*) SulA expression levels. Mean cell brightness is plotted for *recA730* cells grown in the absence and presence of DMSO. Error bars represent standard error of the mean from *n* > 100 cells.

**Fig. S7.**
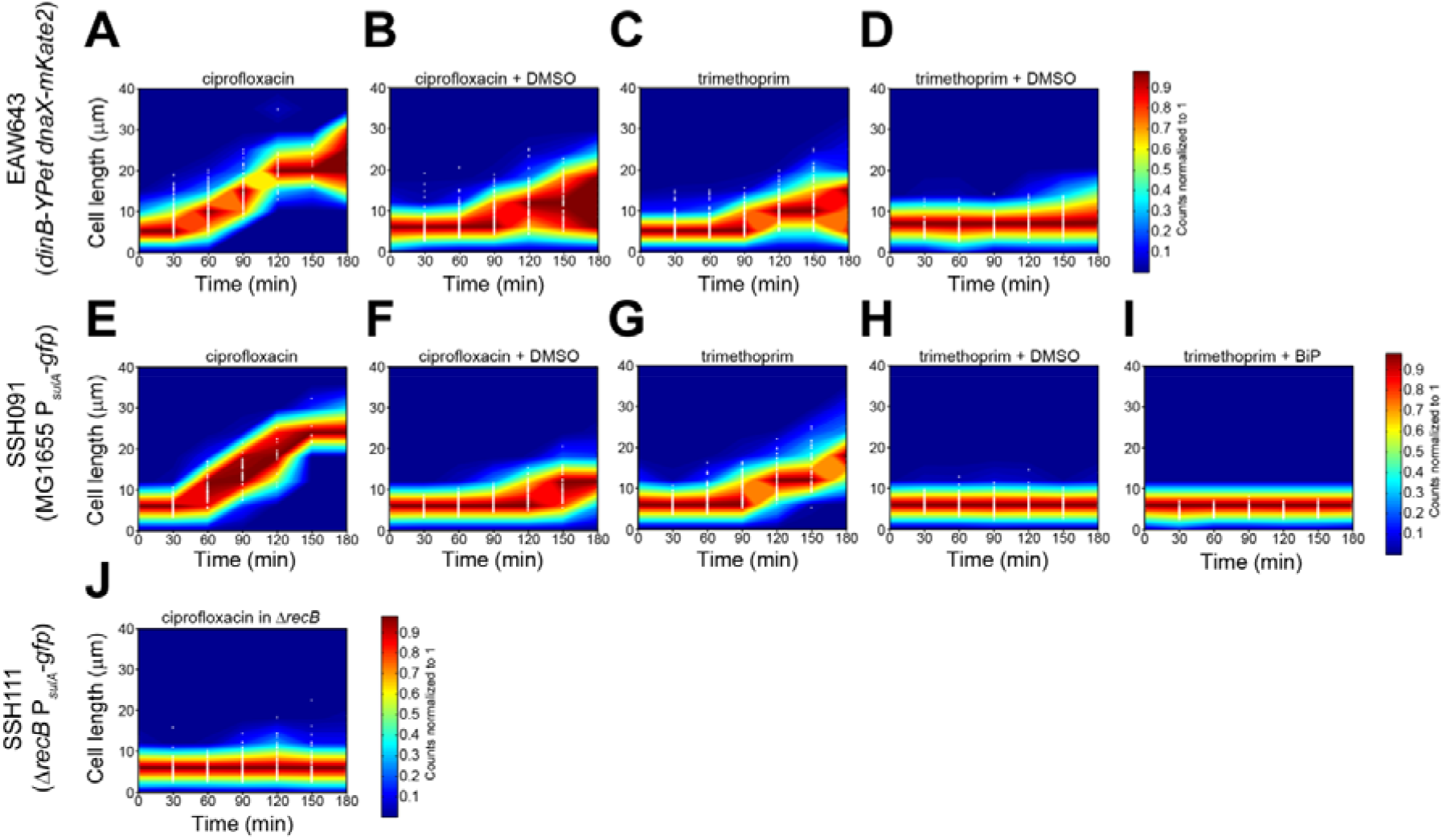
Scatter plots of cell-size from time-lapse imaging. White points indicate individual data-points, while blue-to-red contours indicate frequencies of observations. Blue areas indicate regions of the plot containing few data points; red areas indicate regions containing a large number of data points. Frequencies were normalized at each time-point to the maximum value at that time-point with dark blue = 0 and dark red = 1. We conservatively estimate that >100 cells were used in each measurement. (*A*) EAW643 cells (*dinB-YPet dnaX-mKate2*) treated with ciprofloxacin-alone. (*B*) EAW643 cells (*dinB-YPet dnaX-mKate2*) treated with ciprofloxacin-DMSO. (*C*) EAW643 cells (*dinB-YPet dnaX-mKate2*) treated with trimethoprim-alone. (*D*) EAW643 cells (*dinB-YPet dnaX-mKate2*) treated with trimethoprim-DMSO. (*E*) SSH091 cells (MG1655 P*_sulA_-gfp*) treated with ciprofloxacin-alone. (*F*) SSH091 cells (MG1655 P*_sulA_-gfp*) treated with ciprofloxacin-DMSO. (*G*) SSH091 cells (MG1655 P*_sulA_-gfp*) treated with trimethoprim-alone. (*H*) SSH091 cells (MG1655 P*_sulA_-gfp*) treated with trimethoprim-DMSO. (*I*) SSH091 cells (MG1655 P*_sulA_-gfp*) treated with trimethoprim-BiP. (*J*) SSH111 cells (Δ*recB* P*_sulA_-gfp*) treated with ciprofloxacin.

**Fig. S8.**
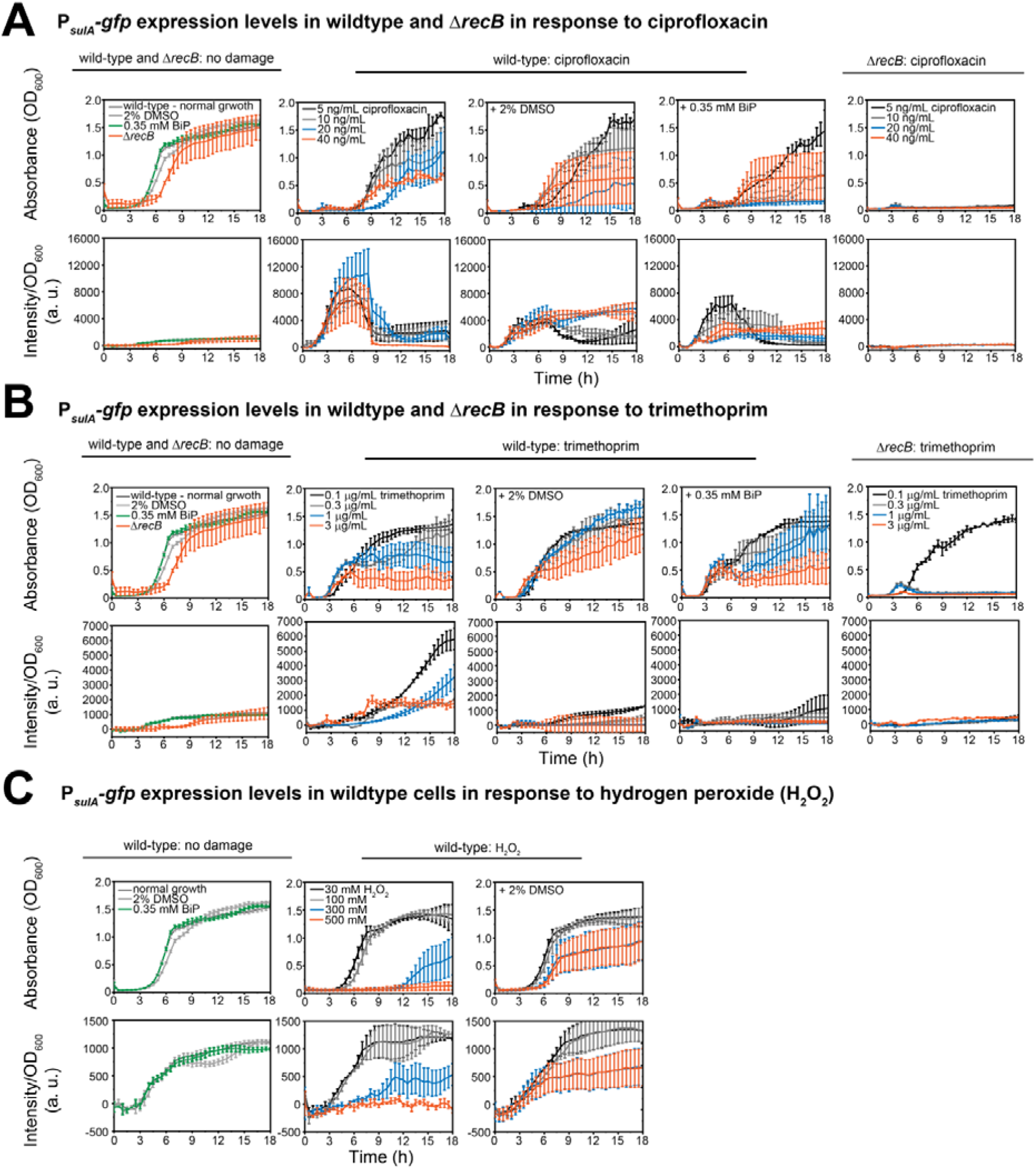
P*_sulA_-gfp* expression levels in wild-type and Δ*recB* cells. For each strain, 10^4^ – 10^6^ cells were added to each well at the beginning of the experiment. Measurements of absorbance (OD_600_) and fluorescence intensity (a.u.) were carried out every 30 min over 18 h. For (A)-(C): upper row shows absorbance (OD_600_) and bottom row illustrates intensity values/ OD_600_, consistent with expression levels. Error bars represent standard error of the mean over three independent biological replicates. (*A*) Comparison of normal growth condition with ciprofloxacin treatment ± ROS mitigator for wild-type cells or Δ*recB.* First column: normal growth conditions (wild-type: dark grey; Δ*recB*: orange), + 2% DMSO (wild-type: grey) or 0.35 mM BiP (wild-type: green); second column: ciprofloxacin treatment of wild-type cells (5 ng/mL: black; 10 ng/mL: grey; 20 ng/mL: blue; 40 ng/mL: orange); third column: ciprofloxacin + 2% DMSO treatment of wild-type cells (same color coding as second column); forth column: ciprofloxacin + 0.35 mM BiP treatment of wild-type cells (same color coding as second column); fifth column: ciprofloxacin treatment of Δ*recB* cells (same color coding as second column). (*B*) Comparison of normal growth condition with trimethoprim treatment ± ROS mitigator for wild-type cells or Δ*recB.* First column: as (A) first column; second column: trimethoprim treatment of wild-type cells (0.1 μg/mL: black; 0.3 μg/mL: grey; 1 μg/mL: blue; 3 μg/mL: orange); third column: trimethoprim + 2% DMSO treatment of wild-type cells (same color coding as second column); forth column: trimethoprim + 0.35 mM BiP treatment of wild-type cells (same color coding as second column); fifth column: trimethoprim treatment of Δ*recB* cells (same color coding as second column). (*C*) Comparison of normal growth condition with hydrogen peroxide (H_2_O_2_) treatment ± ROS mitigator for wild-type cells. First column: as (A) first column; second column: H_2_O_2_ treatment of wild-type cells (30 mM: black; 100 mM: grey; 300 mM: blue; 500 mM: orange); third column: H_2_O_2_ + 2% DMSO treatment of wild-type cells (same color coding as second column).

**Fig. S9.**
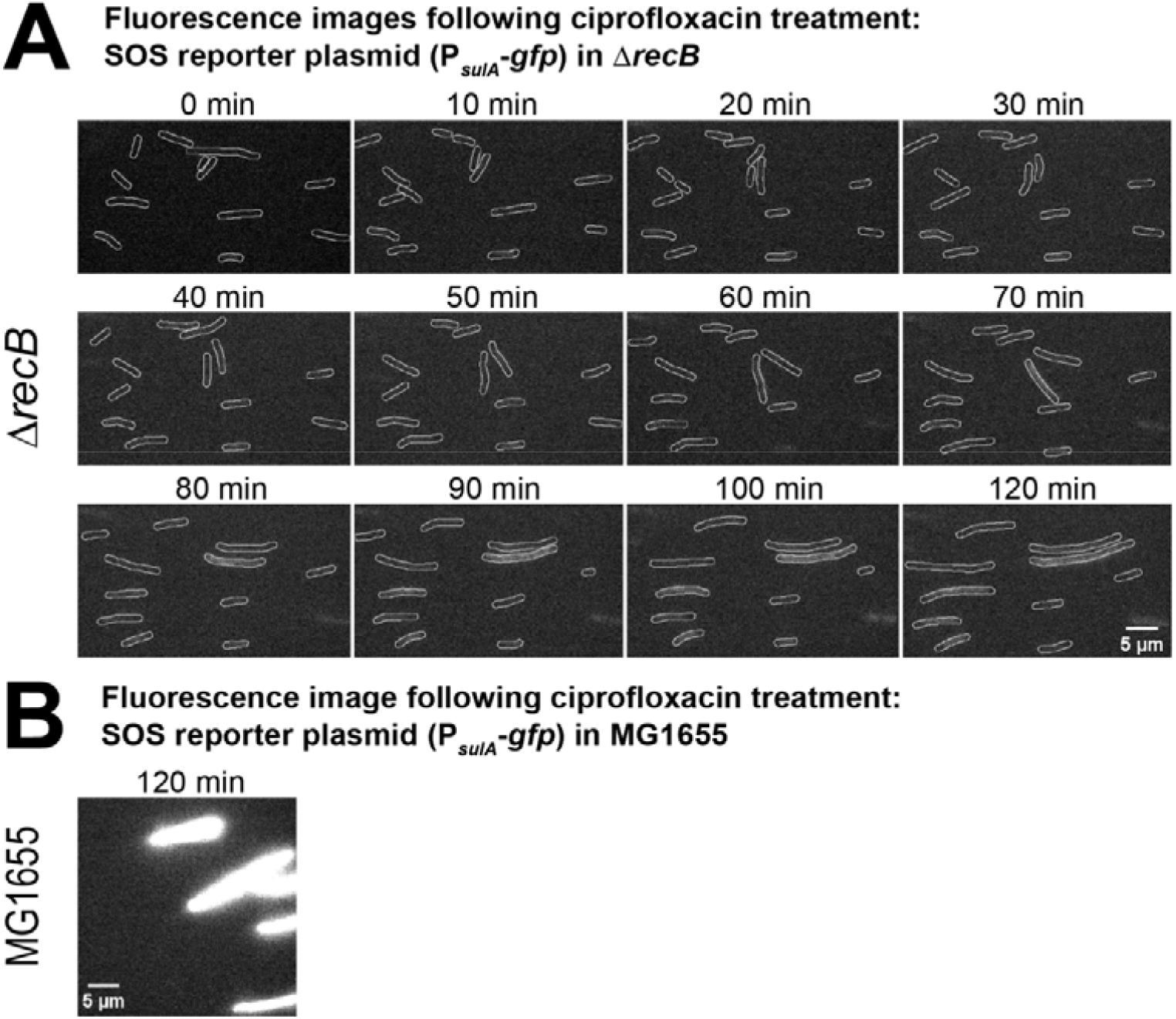
P*_sulA_-gfp* expression levels following ciprofloxacin-alone treatment in ΔrecB vs. MG1655 (wild-type). (*A*) Fluorescence images showing the expression of GFP from a SOS reporter plasmid (P*sulA*-GFP) from 0-110 min at intervals of 10 min and 120 min after ciprofloxacin addition in Δ*recB*. Scale bar represents 5 µm. (*B*) Fluorescence images showing the expression of GFP from a SOS reporter plasmid (P*sulA*-GFP) at 120 min after ciprofloxacin addition in wild-type cells, MG1655. Scale bar represents 5 µm.

**Fig. S10.**
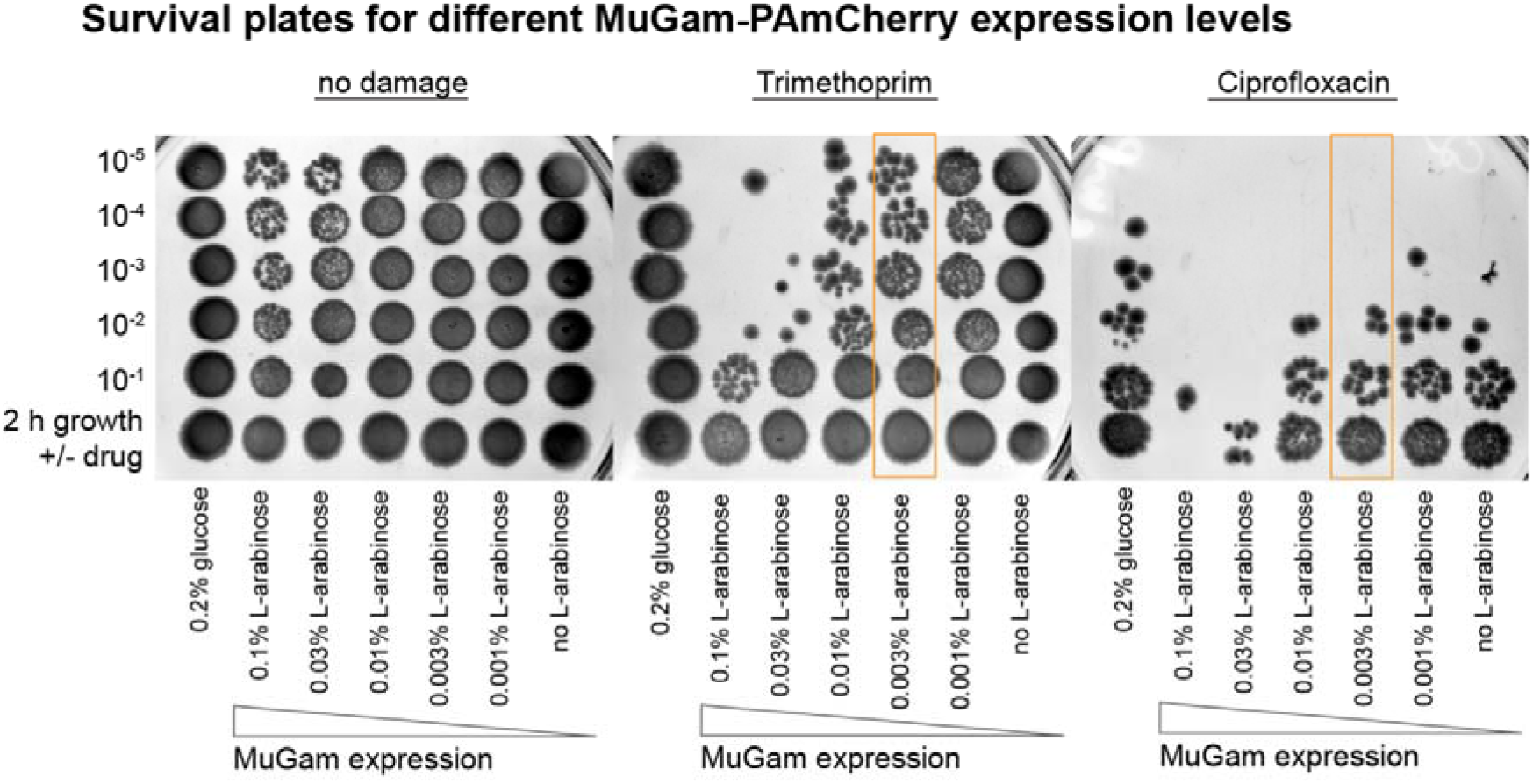
Plate-based survival assays using ciprofloxacin or trimethoprim at different MuGam-PAmCherry expression levels. Cells carrying a pBAD plasmid for MuGam-PAmCherry expression were grown in EZ glycerol in the presence of ampicillin at different L-arabinose concentrations (0, 0.001, 0.003, 0.01, 0.03, 0.1% wt/vol) or in EZ glucose in order to inhibit expression from the pBAD plasmid. These cultures were split in three to perform two survival assays and a ‘no damage’ control. For the survival assays, antibiotic was added to these cultures (30 ng/mL ciprofloxacin or 1 μg/mL trimethoprim), then, cell cultures were grown for 2 h. For the control, cells were grown in the absence of antibiotic for 2 h. After 2 h of growth, cultures were centrifuged and resuspended in glucose or glycerol containing media (x 3) to remove the antibiotic. These cultures were serial diluted in PBS by factor ten down to 10^-5^ and spotted onto LB agar plates containing 100 μg/mL ampicillin. At an L-arabinose concentration of 0.003% (orange box), no drastic decrease in survival was observed in comparison to the sample grown in EZ glucose.

**Fig. S11.**
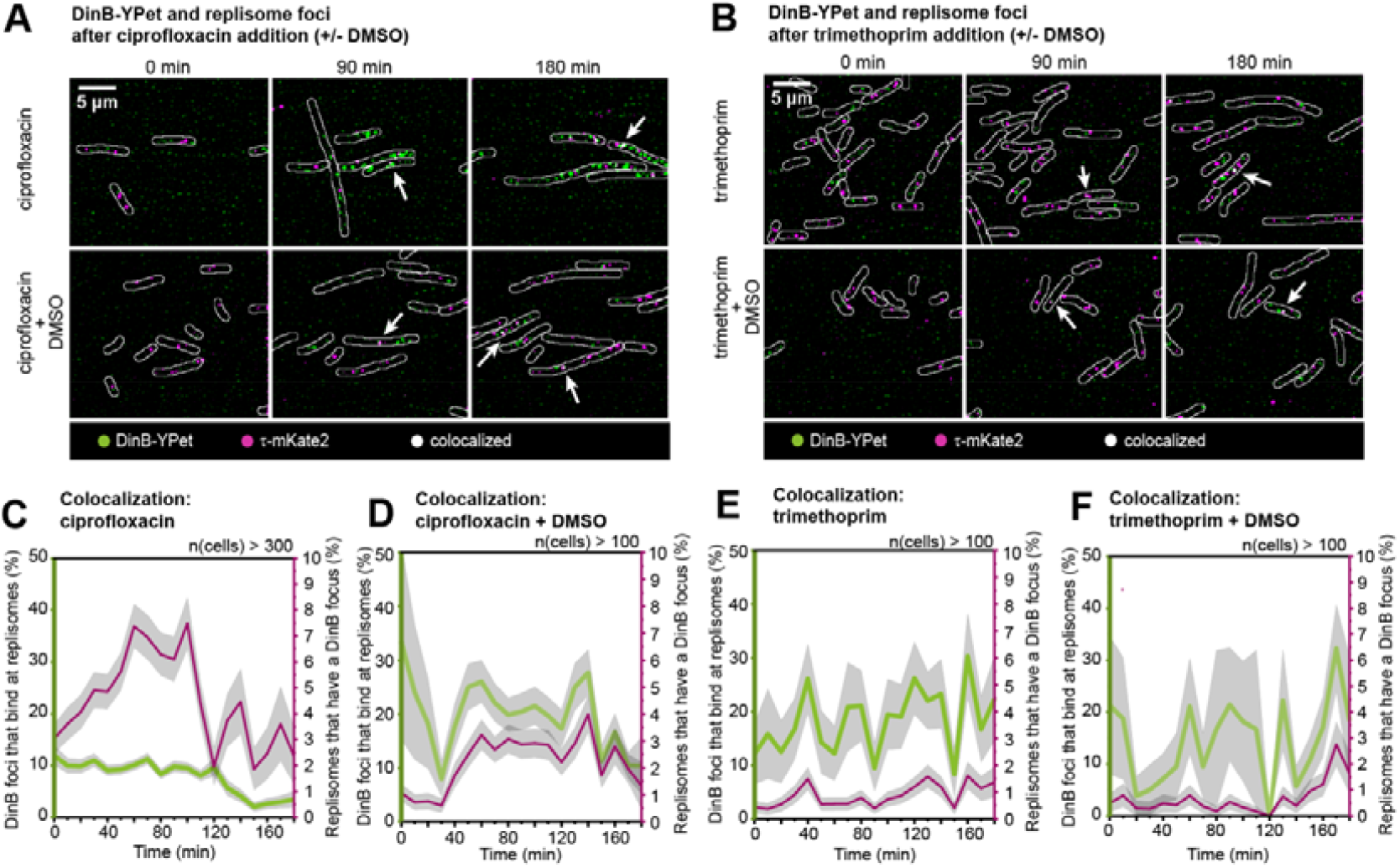
Measuring colocalization of pol IV with replisomes following ciprofloxacin or trimethoprim treatment in the absence or presence of ROS mitigators. (*A*) Merged images showing DinB-YPet (pol IV) foci in green and τ-mKate2 (replisome) foci in magenta at 0, 90 and 180 min (left to right) for ciprofloxacin-alone or ciprofloxacin-DMSO treatment (top to bottom). White arrows indicate colocalization events (white foci). Scale bar represents 5 µm. Merged images showing DinB-YPet (pol IV) foci in green and τ-mKate2 (replisome) foci in magenta at 0, 90 and 180 min (left to right) for trimethoprim-alone or trimethoprim-DMSO treatment (top to bottom). White arrows indicate colocalization events (white foci). Scale bar represents 5 µm. (*C*) Colocalization measurements following ciprofloxacin-alone treatment over 180 min: percentage of pol IV foci that are bound at replisomes (green line), percentage of replisomes that contain a pol IV focus (magenta line). Grey shaded error bands represent the standard error of the mean from six biological replicates together. Measurements are from >300 cells per time point. (*D*) Colocalization measurements following ciprofloxacin-DMSO treatment over 180 min: percentage of pol IV foci that are bound at replisomes (green line), percentage of replisomes that contain a pol IV focus (magenta line). Grey shaded error bands represent the standard error of the mean from four biological replicates together. Measurements are from >100 cells per time point. (*E*) Colocalization measurements following trimethoprim-alone treatment over 180 min: percentage of pol IV foci that are bound at replisomes (green line), percentage of replisomes that contain a pol IV focus (magenta line). Grey shaded error bands represent the standard error of the mean from three biological replicates together. Measurements are from >100 cells per time point. (*F*) Colocalization measurements following trimethoprim-DMSO treatment over 180 min: percentage of pol IV foci that are bound at replisomes (green line), percentage of replisomes that contain a pol IV focus (magenta line). Grey shaded error bands represent the standard error of the mean from three biological replicates together. Measurements are from >100 cells per time point.

**Fig. S12.**
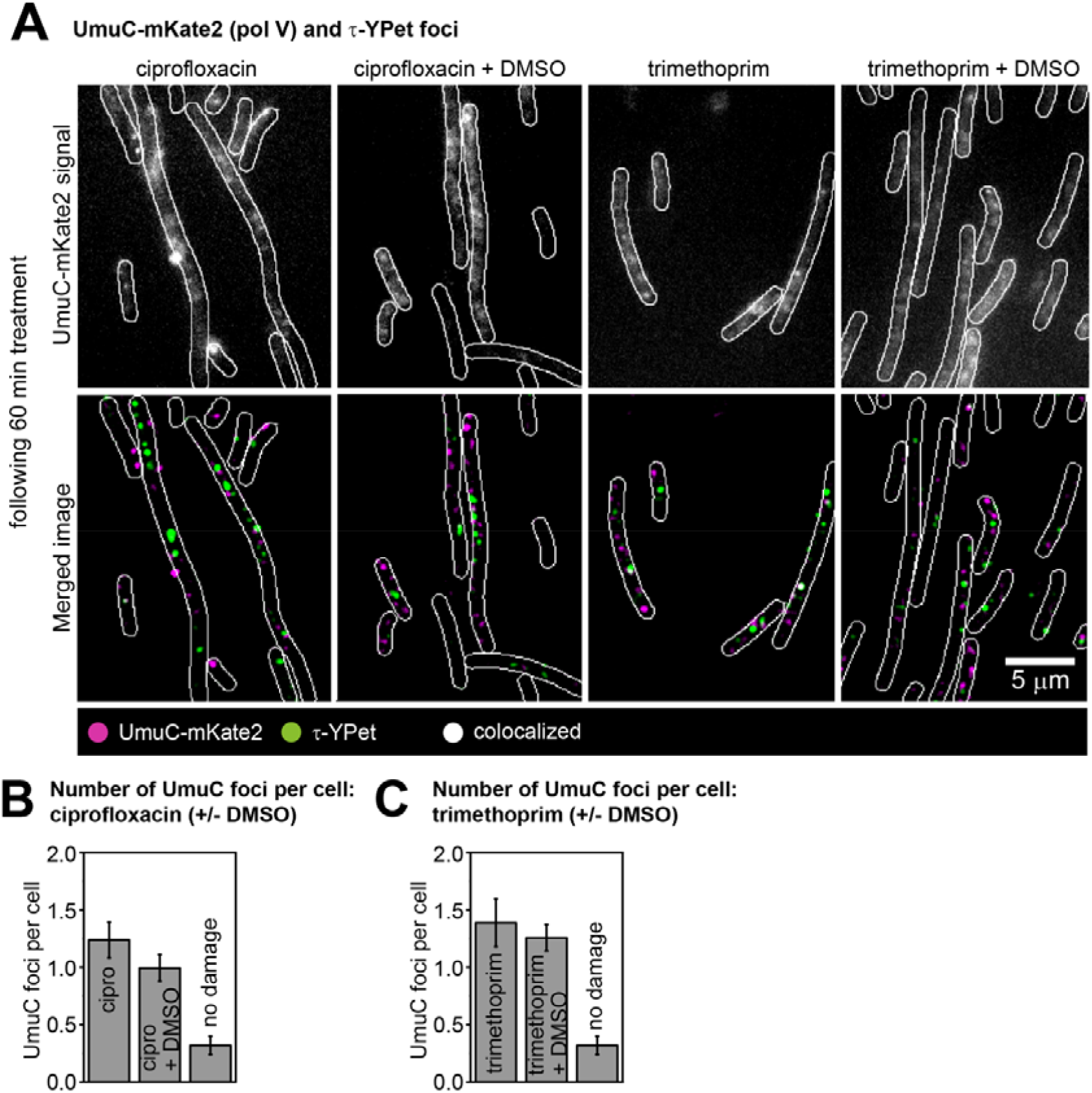
Measuring the number of pol V foci per cell following ciprofloxacin or trimethoprim treatment under normal conditions or ROS-scavenging conditions in *lexA*(Def) cells. (*A*) UmuC-mKate2 activity at replisomes in *lexA*(Def) cells. Cells were treated for 60 min prior to imaging. Upper row: unfiltered image of an average projection showing UmuC-mKate2 foci that last >1 s (from left to right: ciprofloxacin, ciprofloxacin-DMSO, trimethoprim, trimethoprim-DMSO). Bottom row: merged image showing UmuC-mKate2 foci in magenta and τ-YPet foci in green (from left to right: ciprofloxacin, ciprofloxacin-DMSO, trimethoprim, trimethoprim-DMSO). Scale bar represents 5 µm. (*B*) Number of UmuC-mKate2 foci per cell of foci that last > 1 s. Error bars represent standard error of the mean. Number of cells included in analysis: *n*(ciprofloxacin) = 97, *n*(ciprofloxacin-DMSO) = 109, *n*(untreated) = 87. (C) Binding behavior of UmuC-mKate2 at replisomes after ciprofloxacin-alone or ciprofloxacin-DMSO treatment. Mean average autocorrelation function (ciprofloxacin-alone: dark grey line, ciprofloxacin-DMSO: light grey line). Error bar represents standard error of the mean. We conservatively estimate that >400 trajectories from >400 replisomes were used in each measurement. (*D*) Number of UmuC-mKate2 foci per cell. Error bars represent standard error of the mean. Number of cells included in analysis: *n*(trimethoprim) = 102, *n*(trimethoprim-DMSO) = 120, *n*(untreated) = 87. (*E*) Binding behavior of UmuC-mKate2 at replisomes after trimethoprim-alone or trimethoprim-DMSO treatment. Mean average autocorrelation function (trimethoprim-alone: magenta line, trimethoprim-DMSO: light magenta line). Error bar represents standard error of the mean. We conservatively estimate that >550 trajectories from >550 replisomes were used in each measurement.

**Fig. S13.**
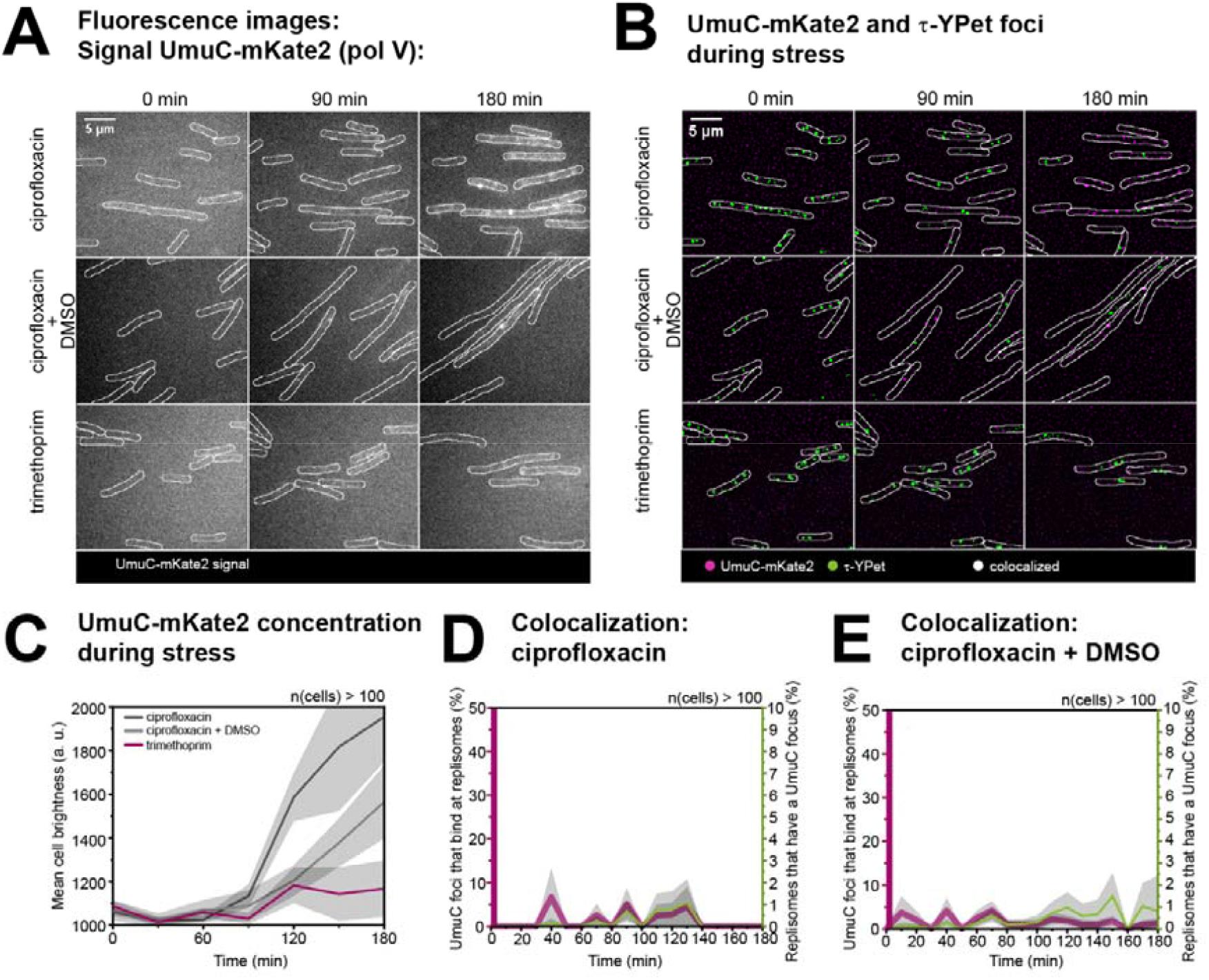
UmuC concentration and activity following ciprofloxacin or trimethoprim treatment under normal conditions or ROS-scavenging conditions. (*A*) Images showing UmuC-mKate2 (pol V) signal at 0, 90 and 180 min (left to right) for ciprofloxacin-alone, ciprofloxacin-DMSO treatment or trimethoprim-alone treatment (top to bottom). Scale bar represents 5 µm. (*B*) Merged images showing UmuC-mKate2 (pol V) foci in magenta and τ-YPet (replisome) foci in magenta at 0, 90 and 180 min (left to right). Colocalized foci would appear as white foci. Scale bar represents 5 µm. (*C*) Concentration of UmuC-mKate2 during stress. Mean cell brightness is plotted against time (ciprofloxacin-alone: dark grey line, ciprofloxacin-DMSO: light grey line, trimethoprim-alone: magenta line). At each time-point, data are derived from >100 cells. Grey shaded error bands represent standard error of the mean. (*D*) Colocalization measurements following ciprofloxacin-alone treatment over 180 min: percentage of UmuC foci that are bound at replisomes (magenta line), percentage of replisomes that contain a UmuC focus (green line). Grey shaded error bands represent the standard error of the mean from three biological replicates together. Measurements are from >100 cells per time point. (*E*) Colocalization measurements following ciprofloxacin-DMSO treatment over 180 min: percentage of UmuC foci that are bound at replisomes (magenta line), percentage of replisomes that contain a UmuC focus (green line). Grey shaded error bands represent the standard error of the mean from three biological replicates together. Measurements are from >100 cells per time point.

**Fig. S14.**
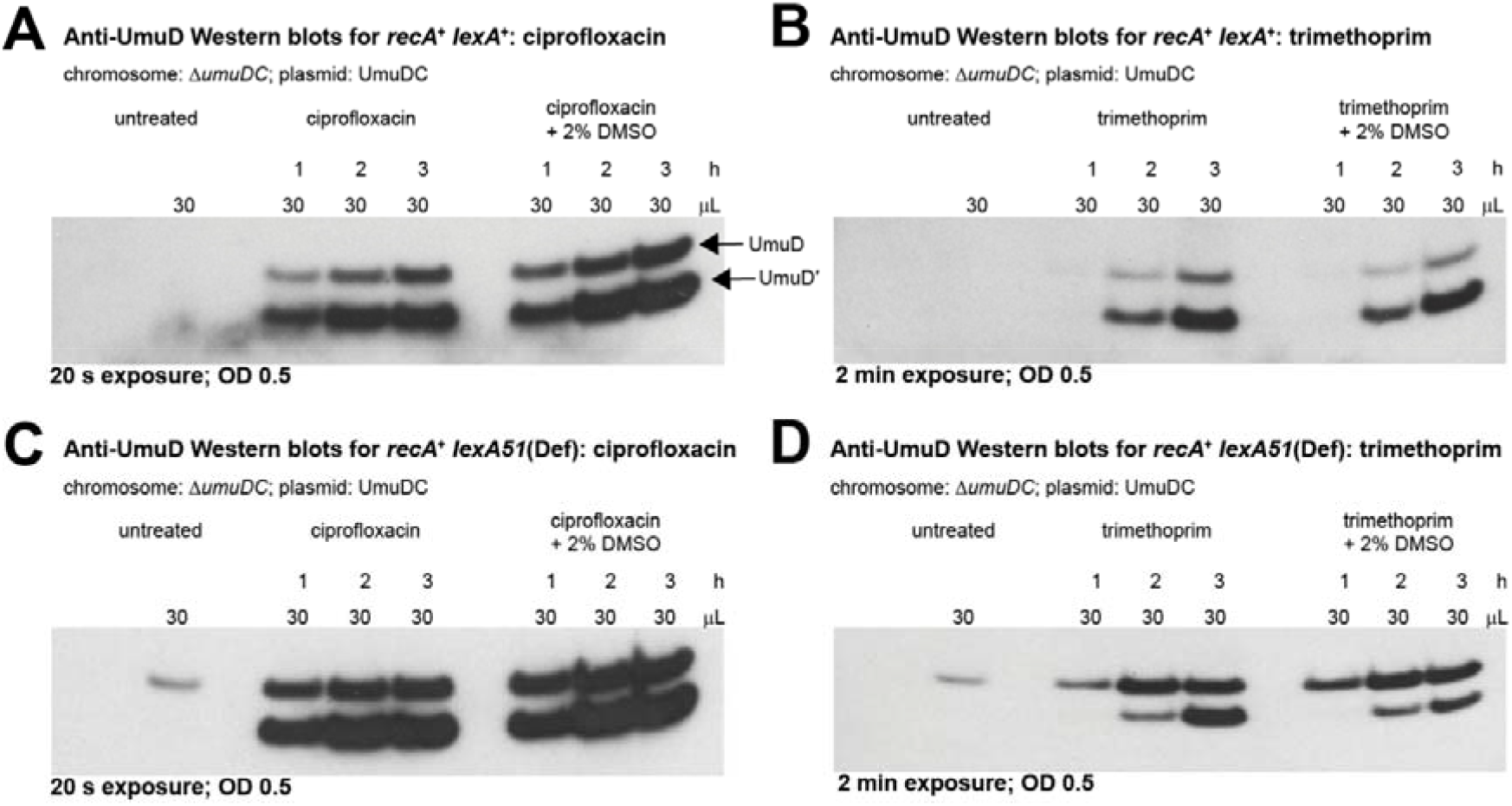
Western blots with anti-UmuD antibodies measuring levels of UmuDD. For each lane, 30 μL of lysate were loaded from cultures at OD_600_ 0.5. All strains used are Δ*umuDC* expressing UmuDC from a low-copy number plasmid (pRW154). After treatment, time points were taken at 1, 2, 3 h. (*A*) Western blot of *recA*^+^ *lexA*^+^ cells (RW120): untreated, treated with ciprofloxacin or ciprofloxacin + 2% DMSO. (*B*) Western blot of *recA*^+^ *lexA*^+^ cells: untreated, treated with trimethoprim or trimethoprim + 2% DMSO. (*C*) Western blot of *recA*^+^ *lexA51*(Def) cells (RW546): untreated, treated with ciprofloxacin or ciprofloxacin + 2% DMSO. (*D*) Western blot of *recA*^+^ *lexA51*(Def) cells: untreated, treated with trimethoprim or trimethoprim + 2% DMSO.

**Movie S1.** Time-lapse movie of Δ*recB* cells carrying P*_sulA_*-*gfp*. Ciprofloxacin (30 ng/mL) was added to the media ay t = 0 min. An image was taken every 10 min over the period of 3 h. Upper movie: bright-field, bottom movie: signal from GFP expression (level of SOS induction).

**Movie S2.** Burst acquisition movies of DinB-YPet in *recB*^+^ cells treated with 1 μg/mL trimethoprim (trimethoprim, ± DMSO) or 30 ng/mL ciprofloxacin (± DMSO), Δ*recB* cells treated with trimethoprim or ciprofloxacin. Movies were recorded 60 min after antibiotic addition. Frames were taken every 0.1 s.

## References

1. X. Zhao, K. Drlica, Reactive oxygen species and the bacterial response to lethal stress. Curr. Opin. Microbiol. 21, 1–6 (2014).

2. H. van Acker, T. Coenye, The role of reactive oxygen species in antibiotic-mediated killing of bacteria. Trends Microbiol. 25, 456–466 (2017).

3. J. A. Imlay, The molecular mechanisms and physiological consequences of oxidative stress: lessons from a model bacterium. Nat. Rev. Microbiol. 11, 443–454 (2013).

4. J. A. Imlay, Where in the world do bacteria experience oxidative stress? Environ. Microbiol. 21, 521–530 (2019).

5. Y. Hong, L. Li, G. Luan, K. Drlica, X. Zhao, Contribution of reactive oxygen species to thymineless death in *Escherichia coli*. Nat. Microbiol. 2, 1667–1675 (2017).

6. A. Liu, L. Tran, E. Becket, K. Lee, L. Chinn et al., Antibiotic sensitivity profiles determined with an *Escherichia coli* gene knockout collection: Generating an antibiotic bar code. Antimicrob. Agents Chemother. 54, 1393–1403 (2010).

7. J. J. Foti, B. Devadoss, J. A. Winkler, J. J. Collins, G. C. Walker, Oxidation of the guinine nucleotide pool underlies cell death by bactericidial antibiotics. Science. 336, 315–319 (2012).

8. D. J. Dwyer, M. A. Kohanski, J. J. Collins, Role of reactive oxygen species in mechanisms of antimicrobial resistance in bacteria. Curr. Opin. Microbiol. 12, 482– 489 (2009).

9. K. Drlica, X. Zhao, DNA gyrase, topoisomerase IV, and the 4-quinolones. Microbiol. Mol. Biol. Rev. 61, 377–392 (1997).

10. J. J. Champoux, DNA topoisomerases: structure, function, and mechanism. Annu. Rev. Biochem. 70, 369–413 (2001).

11. X. Zhao, C. Xu, J. Domagala, K. Drlica, DNA topoisomerase targets of the fluoroquinolones: a strategy for avoiding bacterial resistance. Proc. Natl. Acad. Sci. U.S.A. 94, 13991–13996 (1997).

12. M. Goswami, S. H. Mangoli, N. Jawali, Involvement of reactive oxygen species in the action of ciprofloxacin against *Escherichia coli*. Future Microbiol. 6, 949–954 (2014).

13. X. Giroux, W.-L. Su, M.-F. Bredeche, I. Matic, Maladaptive DNA repair is the ultimate contributor to the death of trimethoprim-treated cells under aerobic and anaerobic conditions. Proc. Natl. Acad. Sci. U.S.A. 114, 11512–11517 (2017).

14. M. A. Kohanski, D. J. Dwyer, B. Hayete, C. A. Lawrence, J. J. Collins, A common mechanism of cellular death induced by bactericidal antibiotics. Cell 130, 797–810 (2007).

15. D. J. Dwyer, J. J. Collins, G. C. Walker, Unraveling the physiological complexities of antibiotic lethality. Annu. Rev. Pharmacol. Toxicol. 55, 313–332 (2015).

16. C. C. Gruber, G. C. Walker, Incomplete base excision repair contributes to cell death from antibiotics and other stresses. DNA Repair. 71, 108–117 (2018).

17. H. Mi, D. Wang, Y. Xue, Z. Zhang, J. Niu et al. Dimethyl sulfoxide protects *Escherichia coli* from rapid antimicrobial-mediated killing. Antimicrob. Agents Chemother. 60, 5054–5058 (2016).

18. K. Drlica, M. Malik, R. J. Kerns, X. Zhao, Quinolone-mediated bacterial death. Antimicrob. Agents Chemother. 52, 385–392 (2008).

19. M. Yamada, T. Nunoshabi, M. Shimizu, P. Grúz, H. Kamiya et al., Involvement of Y-family DNA polymerases in mutagenesis caused by oxidized nucleotides in Escherichia coli. J. Bacteriol. 188, 4992–4995 (2006).

20. J. M. Moore, R. Correa, S. M. Rosenberg, P. J. Hastings, Persistent damaged bases in DNA allow mutagenic break repair in *Escherichia coli*. PLoS Genet. 13, e1006733 (2017).

21. C. Shee, J. L. Gibson, M. C. Darrow, C. Gonzalez, S. M. Rosenberg, Impact of a stress-inducible switch to mutagenic repair of DNA breaks on mutation in *Escherichia coli*. Proc. Natl. Acad. Sci. U.S.A. 108, 13659–13664 (2011).

22. C. Shee, R. Ponder, J. L. Gibson, & S. M. Rosenberg, What limits the efficiency of double-strand break-dependent stress-induced mutation in *Escherichia coli*? J. Mol. Microbiol. Biotechnol. 21, 8–19 (2012).

23. R. G. Ponder, N. C. Fonville, S. M. Rosenberg, A switch from high-fidelity to error-prone DNA double-strand break repair underlies stress-induced mutation. Mol. Cell 19, 791–804 (2005).

24. P. L. Foster, Stress-induced mutagenesis in bacteria. Crit. Rev. Biochem. Mol. Biol. 42, 373–397 (2007).

25. S. M. Rosenberg, C. Shee, R. L. Frisch, P. J. Hastings, Stress-induced mutation *via* DNA breaks in *Escherichia coli*: A molecular mechanism with implications for evolution and medicine. Bioessays 34, 885–892 (2012).

26. V. G. Godoy, D. F. Jarosz, S. M. Simon, A. Abyzov, G. C. Walker, UmuD and RecA directly modulate the mutagenic potential of the Y-family DNA polymerase DinB. Mol. Cell 28, 1058–1070 (2007).

27. S. Mallik, E. M. Popodi, A. J. Hanson, P. L. Foster, Interactions and localization of *Escherichia coli* error-prone DNA Polymerase IV after DNA damage. J. Bacteriol. 197, 2792–2809 (2015).

28. R. T. Pomerantz, I. Kurth, M. F. Goodman, M. O’Donnell, Preferential D-loop extension by a translesion DNA polymerase underlies error-prone recombination. Nat. Struct. Mol. Biol. 20, 748–755 (2013).

29. R. T. Pomerantz, M. F. Goodman, M. E. O’Donnell, DNA polymerases are error-prone at RecA-mediated recombination intermediates. Cell Cycle 12, 2558–2563 (2013).

30. C. Indiani, M. Patel, M. F. Goodman, M. E. O’Donnell, RecA acts as a switch to regulate polymerase occupancy in a moving replication fork. Proc. Natl. Acad. Sci. U.S.A. 110, 5410–5415 (2013).

31. T. M. Cafarelli, T. J. Rands, V. G. Godoy, The DinB·RecA complex of *Escherichia coli* mediates an efficient and high-fidelity response to ubiquitous alkylation lesions. Environ. Mol. Mutagen. Mutagen. 55, 92–102 (2014).

32. S. S. Henrikus et al., UmuD and RecA* modulate the DNA-binding activity of DNA polymerase IV in *Escherichia coli*. bioRXiv:620195 (29 April 2019).

33. T. F. Tashjian, C. Danilowicz, A.-E. Molza, B. H. Nguyen, C. Prévost et al. Residues in the fingers domain of the translesion DNA polymerase DinB enable its unique participation in error-prone double strand break repair. J. Biol. Chem. jbc.RA118.006233 (2019).

34. G. R. Smith, Homologous recombination in prokaryotes: enzymes and controlling sites. Genome 31, 520–527 (1989).

35. L. A. Simmons, J. J. Foti, S. E. Cohen, G. C. Walker, The SOS regulatory network. EcoSal Plus 3, doi:10.1128/ecosalplus.5.4.3 (2008).

36. A. McPartland, L. Green, H. Echols, Control of *recA* gene RNA in *E. coli*: Regulatory and signal genes. Cell 20, 731–737 (1980).

37. K. G. Newmark, E. K. O’Reilley, J. R. Pohlhaus, K. N. Kreuzer, Genetic analysis of the requirements for SOS indcution by nalidixic acid in *Escherichia coli*. Gene 356, 69–76 (2005).

38. J. Courcelle, A. Khodursky, B. Peter, P. O. Brown, P. C. Hanawalt, Comparative gene expression profiles following UV exposure in wild-type and SOS-deficient. Genetics 158, 41–64 (2001).

39. S. S. Henrikus, E. A. Wood, J. P. McDonald, M. M. Cox, R. Woodgate et al., DNA polymerase IV primarily operates outside of DNA replication forks in *Escherichia coli*. PLoS Genet. 14, e1007161 (2018).

40. Imlay, J. A. Transcription factors that defend bacteria against reactive oxygen species. Annu. Rev. Microbiol. 69, 93–108 (2015).

41. S. W. Seo, D. Kim, R. Szubin, B. O. Palsson, Genome-wide reconstruction of OxyR and SoxRS transcriptional regulatory networks under oxidative stress in *Escherichia coli* K-12 MG1655. Cell Rep. 12, 1289–1299 (2015).

42. G. Storz, L. A. Tartaglia, B. N. Ames, The OxyR regulon. Antonie van Leeuwenhoek, Int. J. Gen. Mol. Microbiol. 58, 157–161 (1990).

43. I. L. Jung, I. G. Kim, Transcription of *ahpC*, *katG*, and *katE* genes in *Escherichia coli* is regulated by polyamines: Polyamine-deficient mutant sensitive to H_2_O_2_-induced oxidative damage. Biochem. Biophys. Res. Commun. 301, 915–922 (2003).

44. J. L. Lavrrar, C. A. Christoffersen, M. A. McIntosh, Fur-DNA interactions at the bidirectional *fepDGC-entS* promoter region in *Escherichia coli*. J. Mol. Biol. 322, 983– 995 (2002).

45. E. A. Ronayne, Y. C. S. Wan, B. A. Boudreau, R. Landick, M. M. Cox, P1 Ref endonuclease: a molecular mechanism for phage-enhanced antibiotic lethality. PLoS Genet. 12, e1005797 (2016).

46. M. Goswami, S. H. Mangoli, N. Jawali, Involvement of reactive oxygen species in the action of ciprofloxacin against *Escherichia coli*. Antimicrob. Agents Chemother. 50, 949–954 (2006).

47. A. Robinson, J. P. McDonald, V. E. A. Caldas, M. Patel, E. A. Wood et al., Regulation of mutagenic DNA polymerase V activation in space and time. PLoS Genet. 11, e1005482 (2015).

48. G.-W. Li, X. Sunney Xie. Central dogma at the single-molecule level of living cells. Nature 475, 308–315 (2011).

49. A. Zaslaver, A. Bren, M. Ronen, S. Itzkovitz, I. Kikoin et al., A comprehensive library of fluorescent transcriptional reporters for *Escherichia coli*. Nat. Methods 3, 623–628 (2006).

50. A. Rodriguez-Rosado et al., Reactive oxygen species are major contributors to SOS-mediated mutagenesis induced by fluoroquinolones. bioRXiv:428961 (27 September 2018).

51. L. L. Shen, A. G. Pernet, Mechanism of inhibition of DNA gyrase by analogues of nalidixic acid: The target of the drugs is DNA. Proc. Natl. Acad. Sci. U.S.A. 82, 307–311 (1985).

52. T. Dorr, K. Lewis, M. Vulic, SOS response induces persistence to fluoroquinolones in *Escherichia coli*. PLoS Genet. 5, e1000760 (2009).

53. C. Shee, B. D. Cox, F. Gu, E. M. Luengas, M. C. Joshi et al., Engineered proteins detect spontaneous DNA breakage in human and bacterial cells. Elife 2, 1–25 (2013).

54. H. Ghodke, B. Paudel, J. Lewis, S. Jergic, K. Gopal et al., Spatial and temporal organization of RecA in the *Escherichia coli* DNA-damage response. Elife 8, e42761 (2019).

55. J. P. Pribis, L. García-Villada, Y. Zhai, O. Lewin-Epstein, A. Wang et al. Gamblers: an antibiotic-induced evolvable cell subpopulation differentiated by reactive-oxygen-induced general stress response. Mol. Cell 74, 785–800 (2019).

56. D. G. Ennis, B. Fisher, S. Edmiston, D. W. Mount, Dual role for *Escherichia coli* RecA protein in SOS mutagenesis. Proc. Natl. Acad. Sci. U.S.A. 82, 3325–3329 (1985).

57. M. Snyder, K. Drlica, DNA gyrase on the bacterial chromosome: DNA cleavage induced by oxolinic acid. J. Mol. Biol. 131, 287–302 (1979).

58. W. H. Deitz, T. M. Cook, W. A. Goss, Mechanism of action of nalidixic acid on *Escherichia coli* III. Conditions required for lethality. J. Bacteriol. 91, 768–773 (1966).

59. M. Tang, X. Shen, E. G. Frank, M. O’Donnell, R. Woodgate et al., UmuDD_(2)_C is an error-prone DNA polymerase, *Escherichia coli* pol V. Proc. Natl. Acad. Sci. U.S.A. 96, 8919–8924 (1999).

60. R. T. Cirz, J. K. Chin, D. R. Andes, V. de Crécy-Lagard, W. A. Craig et al., Inhibition of mutation and combating the evolution of antibiotic resistance. PLoS Biol. 3, e176 (2005).

61. A. Sakai, M. Nakanishi, K. Yoshiyama, H. Maki, Impact of reactive oxygen species on spontaneous mutagenesis in *Escherichia coli*. Genes to Cells 11, 767–778 (2006).

62. R. Gleckman, N. Blagg, D. W. Joubert, Trimethoprim: Mechanisms of action, antimicrobial activity, bacterial resistance, pharmacokinetics, adverse reactions, and therapeutic indications. Pharmacotherapy 1, 14–19 (1981).

63. B. Birdsall, G. C. Roberts, J. Feeney, J. Dann, A. Burgen, Trimethoprim binding to bacterial and mammalian dihydrofolate reductase: a comparison by proton and carbon-13 nuclear magnetic resonance. Biochemistry 22, 5597–5604 (1983).

64. I. Vlašić, I. Ivančić-Baće, M. Imešek, B. Mihaljević, K. Brčić-Kostić, RecJ nuclease is required for SOS induction after introduction of a double-strand break in a RecA loading deficient *recB* mutant of *Escherichia coli*. Biochimie 90, 1347–1355 (2008).

65. I. Vlašić, A. Šimatović, K. Brčić-Kostić, Genetic requirements for high constitutive SOS expression in *recA730* mutants of *Escherichia coli*. J. Bacteriol. 193, 4643–4651 (2011).

66. K. Morimatsu, S. C. Kowalczykowski, RecFOR proteins load RecA protein onto gapped DNA to accelerate DNA strand exchange: A universal step of recombinational repair. Mol. Cell 11, 1337–1347 (2003).

67. A. Sakai, M. M. Cox, RecFOR and RecOR as distinct RecA loading pathways. J. Bio 284, 3264–3272 (2009).

68. C. Lesterlin, G. Ball, L. Schermelleh, D. J. Sherratt, RecA bundles mediate homology pairing between distant sisters during DNA break repair. Nature 506, 249–253 (2014).

69. E. G. Frank, D. G. Ennis, M. Gonzalez, A. S. Levine, R. Woodgate, Regulation of SOS mutagenesis by proteolysis. Proc. Natl. Acad. Sci. U.S.A. 93, 10291–6 (1996).

70. L. C. Huang, E. A. Wood, M. M. Cox, Convenient and reversible site-specific targeting of exogenous DNA into a bacterial chromosome by use of the FLP recombinase: The FLIRT system. J. Bacteriol. 179, 6076–6083 (1997).

71. D. G. Ennis, S. K. Amunsden, G. R. Smith, Genetic functions promoting homologous recombination in *Escherichia coli*: A study of inversions in phage λ. Genetics 115, 11– 24 (1987).

72. E. G. Frank, J. Hauser, A. S. Levine, R. Woodgate, Targeting the UmuD, UmuDD, and MucAD mutagenesis proteins to DNA by RecA protein. Proc. Natl. Acad. Sci. U.S.A. 90, 8169–8173 (1993).

73. D. R. Harris, S. V. Pollock, E. A. Wood, R. J. Goiffon, A. J. Klingele et al., Directed evolution of ionizing radiation resistance in *Escherichia coli*. J. Bacteriol. 191, 5240– 5252 (2009).

74. F. R. Blattner, G. Plunkett III, C. A. Bloch, N. T. Perna, V. Burland et al., The complete genome sequence of *Escherichia coli* K-12. Science. 277, 1453–1474 (1997).

75. C. Ho, O. I. Kulaeva, A. S. Levine, R. Woodgate, A rapid method for cloning mutagenic DNA repair genes: Isolation of *umu*-complementing genes from multidrug resistance plasmids R391, R446b, and R471a. J. Bacteriol. 175, 5411–5419 (1993).

76. S. Rangarajan, R. Woodgate, M. F. Goodman, Replication restart in UV-irradiated *Escherichia coli* involving pols II, III, V, PriA, RecA and RecFOR proteins. Mol. Microbiol. 43, 617–628 (2002).

77. C. A. Schneider, W. S. Rasband, K. W. Eliceiri, NIH Image to ImageJ: 25 years of image analysis. Nat. Methods 9, 671–675 (2012).

78. O. Sliusarenko, J. Heinritz, T. Emonet, C. Jacobs-Wagner, High-throughput, subpixel-precision analysis of bacterial morphogenesis and intracellular spatio-temporal dynamics. Mol. Microbiol. 80, 612–627 (2012).

